# A lipid-binding protein in black-legged tick saliva selectively recognizes *Borrelia burgdorferi* lipids

**DOI:** 10.64898/2026.05.04.722819

**Authors:** Weirui Olivia Shi, Taylor MacMackin-Ingle, Mia Wong Perez, Wendell P. Griffith, Liao Y. Chen, J. Seshu, Robert Renthal

## Abstract

A proteomic analysis of *Ixodes scapularis* nymph saliva identified 252 proteins, including six tubular lipid-binding proteins (TULIPs). Comparing nymphs fed on mice that were uninfected or infected with *Borrelia burgdorferi*, twelve salivary proteins showed significant differences in the amounts detected, including XP_040079658.2, which we refer to as TULIP2. Considering the known immunity-related functions of some TULIPs, we expressed and purified TULIP2 from *Escherichia coli* and analyzed its interaction with *B. burgdorferi* lipids. The purification of TULIP2 from *E. coli* presented many obstacles, due to insolubility, which is consistent with previous reports from studies of other TULIP family members. The binding results showed specificity for *B. burgdorferi* lipids, with evidence for cholesteryl β-galactoside as a major binding target. Molecular modeling of TULIP2 did not show any strong lipid binding sites. We used molecular dynamics simulation of TULIP2 to explore its conformational landscape by thermal unfolding. The earliest unfolding intermediate opened a hydrophobic pocket to which cholesteryl β-galactoside was predicted to bind strongly. We propose that a specific lipid bilayer interaction with TULIP2 triggers the opening of the ligand-binding site.

## 1. Introduction

*Ixodes scapularis* is a vector of multiple disease pathogens, including *Borrelia burgdorferi* s.l., *Anaplasma phagocytophilum*, *Borrelia miyamotoi*, *Babesia* sp., Powassan virus, and *Ehrlichia muris eauclarensis*^1,2^. When a tick is attached to a vertebrate host and taking a bloodmeal, it injects saliva at the host bite site. As a slow-feeding tick, *I. scapularis* takes a few days to finish a bloodmeal, so it secretes compounds that secure the attachment, maintain blood flow at the bite site, modulate the host’s immune defense, and suppress pain or itch ^3^. Because of the risk of taking up pathogens in a bloodmeal, ticks have several weapons to defend against infection. Nevertheless, each species has a niche of a few pathogens that it carries. Ticks have three major immune pathways, namely the Toll pathway, the immune deficiency (IMD) pathway, and RNAi pathways ^4,5^. Other components of the *I. scapularis* immune system include antimicrobial peptides, agglutination proteins, phagocytosis, coagulation, or proteolysis ^5–7^.

In view of the role of pathogen lipid detection in *I. scapularis* immunity ^8,9^, we were intrigued when we observed six paralogous tubular lipid-binding proteins (TULIPs) expressed in *I. scapularis* saliva. We considered the possibility that one or more of these proteins could be involved in signaling the presence of pathogens to the tick immune system. TULIPs belong to a large superfamily of eukaryotic proteins, as mediators of lipid-sensing and transport, including some TULIP family members that are involved in immune responses ^10,11^. Three-dimensional structures of some vertebrate and insect TULIPs show internal tube-like cavities ^12–15^, to which lipids are known to bind, sometimes with high selectivity. However, structures of other TULIPs have much smaller internal cavities, even in cases where lipid binding is known to occur ^16^, suggesting that a closed-to-open conformational transition may accompany ligand binding.

Of the six TULIPs we detected in *I. scapularis* saliva, only TULIP2 showed significant differences between levels in *I. scapularis* nymphs lacking or harboring *B. burgdorferi*. Therefore, in this report, we have examined the lipid-binding properties of TULIP2 as a possible participant in the *I. scapularis* immune response to *B. burgdorferi*.

## 2. Materials and methods

### 2.1 Biological materials and sources

*Ixodes scapularis* larvae were purchased from the tick rearing facility at Oklahoma State University (Stillwater, OK) and incubated in a chamber at 22°C with 99% humidity and a 12-hour light / 12-hour dark cycle. *Borrelia burgdorferi* B31-A3 was grown in liquid Barbour-Stoenner-Kelly (BSK II) media supplemented with 6% heat inactivated bovine serum at pH 7.6, 32 °C.

### 2.2 Animals and ethics statement

All animals used in this study were housed in the facilities of Laboratory Animal Resources Center (LARC), which is an AAALAC International Accredited Unit at The University of Texas at San Antonio (UTSA). Six-to eight-week-old female C3H/HeN and BALB/c mice (Charles River Laboratories, Wilmington, MA) were used in this study. All animal experiments were conducted following standard guidelines for housing and care of laboratory animals and in accordance with protocol approved by the Institutional Animal Care and Use Committee of UTSA. Based on these guidelines, general condition and behavior of the animals were monitored by trained laboratory and LARC staff daily and methods to minimized pain and discomfort were adopted as needed in this study. All animals were euthanized using CO_2_ inhalation followed by cardiac puncture as the second method of euthanasia.

Mouse infections: A low passage infectious clonal isolate of *B. burgdorferi* strain B31-A3 (Bb) propagated in BSKII media supplemented with 6% normal rabbit serum at 32°C/pH7.6 to a density of 1to2 ×10^7^ Bb/ml were washed in 50% normal rabbit serum in HBSS thrice (with centrifugation at 8000xg in a microfuge) and resuspended at 1×10^6^ *Bb*/ml. A volume of one hundred microliters of the sample is injected intradermally ^17–19^. Three to 5 days post-challenge, naïve Ixodes scapularis larvae hatched from egg masses (purchased from the tick rearing facility at Oklahoma State University (Stillwater, OK) were allowed to feed by placing them on capsules glued to shaved dorsal surface of mouse. Larvae were fed to repletion on naïve and infected C3H/HeN mice and were collected and incubated in a chamber at 22°C with 99% humidity and a 12-hour light / 12-hour dark cycles till they molted to nymphs. Nymphs were allowed to feed on naïve mice to repletion, and fed nymphs were used to obtain the saliva, as described below ^17–19^.

### 2.3 Collection of tick saliva

A volume of 1 to 2ul of 2% pilocarpine hydrochloride, prepared in phosphate buffered saline pH 7.4, was injected on the ventral side of the lower right coxa with a Hamilton syringe to induce tick salivation (Hamilton Company, Reno, NV, USA) (Triloni et al 2017). Tick saliva was lyophilized and stored at −80°C. Saliva from 4 infected and 5 uninfected nymphs were collected and processed as described below.

### 2.4 TULIP2 plasmids

The DNA sequence coding for TULIP2 was obtained as a gBlock from IDT (Coralville, IA). Initially, TULIP2 was expressed with a C-terminal 6× His tag from a pET23a vector, with NdeI and XhoI, respectively, the 5’ and 3’ insertion sites flanking TULIP2. Another expression vector encoding an N-terminal maltose-binding protein (MBP) tag was constructed by using BamHI and SalI sites to insert TULIP2 DNA into a modified pMAL-c2x vector that had the thrombin cleavage site replaced by a TEV protease site. The enzyme-digested TULIP2-encoding DNA was purified by ethanol precipitation. The purified empty plasmid and digested gene were combined at a ratio of 1:3 and ligated with T4 DNA ligase (NEB) at room temperature for two hours. The ligation product was delivered into Rosetta 2 competent cells (ThermoFisher) by heat shock. Three colonies were picked out after transformation. The plasmid insert sequence was confirmed by Sanger sequencing (Eurofins).

### 2.5 Expression and purification of recombinant TULIP2

Rosetta 2 (*E. coli* DE3) competent cells (Novagen) were transformed by the expression plasmids using heat shock. After transformation, cells were recovered for one hour in Luria-Bertani (LB) broth at 37 °C. The cells were spread on an LB-agar plate containing 20 µg/ml chloramphenicol and 100 µg/ml carbenicillin and cultured overnight. Single colonies were isolated and expanded in 6 ml LB medium supplemented with 100 mg/L carbenicillin and 20 mg/L chloramphenicol. After 6-hour incubation at 37 °C with shaking, the culture was diluted 1000-fold into fresh LB medium with antibiotics and cultured overnight. On the next morning, 0.2 mM isopropyl β-D-1-thiogalactopyranoside (IPTG) was introduced to the culture and the incubation temperature was lowered to 30 °C. After six hours of IPTG induction, cells were harvested by centrifugation at 5000 × g, 4 °C for 20 min. Cells were washed twice in ice-cold phosphate-buffered saline (PBS).

For purification of the His-tagged TULIP2, the bacteria were resuspended in PBS and lysed in B-Per bacterial protein extraction reagent (ThermoFisher). The cell lysate was centrifuged at maximum speed in a microcentrifuge (Eppendorf 5814R) at 4 °C for 20 min. The recombinant protein TULIP2-6×His was recovered in inclusion bodies. The inclusion body pellet was washed in 1 % Triton X-100 and dissolved in 5 M guanidinium hydrochloride (Gdn). Purification was performed using a HisPur Ni-NTA spin column (ThermoFisher) according to the manufacturer’s instructions, and all buffers used were supplemented with 5 M Gdn.

For purification of MBP-TULIP2, *E. coli* cells were harvested, washed, resuspended in PBS, and lysed by a French pressure cell (ThermoFisher). The cell lysate was centrifuged at maximum speed in a microcentrifuge (Eppendorf 5814R) at 4 °C for 20 min. The supernatant was separated for protein purification through an MBPTrap HP maltose affinity column (Cytiva) following manufacturer’s recommended Tris buffers and procedures. For MBP removal, the elution fractions containing MBP-TULIP2 were digested with TEV protease (NEB) at 4 °C overnight.

### 2.6 Mouse TULIP2 antibody

TULIP2-6×His was expressed in *E. coli* and purified using Ni-NTA column and the His-tag cleaved with TEV protease. Recombinant TULIP2 without a His-tag was then emulsified in equal volume of TiterMax gold adjuvant (Sigma), and used to immunize six to eight weeks old BALB/c mice at 50 micrograms/mouse (day 0). Boosters were administered at day 14 and 21, and serum was collected on day 28 ^19,20^.

### 2.7 *Borrelia burgdorferi* total lipid extraction

Total lipids were extracted from *B. burgdorferi* B31cultured in BSK medium using phase separation in chloroform-methanol-water ^21^. A pellet containing approximately 10^8^ *B. burgdorferi* cells was resuspended in 0.8 ml water and added to 3.75 ml organic solvent (methanol : chloroform 2 : 1) in a capped teflon tube. The cells were rocked at room temperature for 1 hour. The mixture was centrifuged, and the same amount of methanol/chloroform/water was added to the pellet for a second extraction. After centrifugation, 2.5 ml chloroform and 2.5 ml water were added to the combined extract for phase separation. The chloroform phase was withdrawn and evaporated under a nitrogen stream to concentrate the lipids by three-fold. The concentration of phospholipids in the chloroform phase was measured by organic phosphate assay ^22^. The total lipid content was estimated based on the assumption that phospholipids represent 20 % of the total lipids in *B. burgdorferi* ^23^.

### 2.8 Lipid dot blot

Control samples for the lipid dot blots were selected to contain functional groups representative of the known composition of *B. burgdorferi* membrane lipids ^23–25^: 1-palmitoyl-2-oleoylphosphatidylcholine (Avanti), cholesteryl stearate (Sigma), 1,2-dipalmitoylglycerol (Avanti), and ⍺-galactosyl ceramide (Avanti). They were dissolved in chloroform or chloroform/methanol to make a 3-6 mg/ml stock solution. Lipid samples were repeatedly spotted onto nitrocellulose membranes, 1 µl at a time, to reach a final amount in the range of ∼1-10 nanomole in each dot ^26^. The lipid dots were air-dried before the membrane was blocked in 3 % BSA. The blocked membrane was incubated with TULIP2 (7-14 μM) in blocking buffer at 4 °C overnight. The blocked membrane was rinsed three times in Tris-buffered saline with Tween-20 (TBST: 25 mM Tris, 120 mM NaCl, pH 7.5, with 0.1 % Tween-20), 10 min. each. Mouse anti-TULIP2 antiserum (1: 5000) and horseradish peroxidase-conjugated goat anti-mouse IgG (Invitrogen) (1 : 5000) were applied sequentially by one-hour incubation at room temperature, with washing as described above following each antibody. Chemiluminescence detection and imaging was performed in a ChemiDoc imager (Bio-Rad). Dot intensity was measured using ImageJ software (NIH), and intensities from ∼0.7 nmol *B. burgdorferi* lipids were used to normalize intensities from separate blots. Pairs of different lipids were compared by a one-tailed t-test.

### 2.9 Immunoblot Analysis

Proteins were resolved by sodium dodecyl sulfate polyacrylamide gel electrophoresis (SDS-PAGE) on precast Bis-Tris 4-12 % gradient gels (Invitrogen, ThermoFisher), run in 2-(N-morpholino)ethanesulfonic acid (MES) buffer, and transferred to a nitrocellulose membrane using a Trans-Blot electrophoretic transfer cell (BioRad). The membrane was blocked in 3 % BSA. Mouse anti-TULIP2 antiserum or mouse anti-MBP (Sigma-Aldrich) was diluted in blocking buffer at 1:5000 and applied to the blot at 4 °C overnight. The secondary antibodies were HRP-conjugated goat anti-mouse IgG (Invitrogen) diluted 5000-fold in TBST. After each antibody incubation, unbound antibodies were washed in TBST. Chemiluminescence was recorded in a ChemiDoc imager (Bio-Rad).

### 2.10 Protein analysis

Saliva extracts were added directly to NuPAGE LDS sample buffer (ThermoFisher) and separated on short (∼5 cm) Bis-Tris 4-12% polyacrylamide electrophoresis gels (ThermoFisher), run in MES buffer, pH 7.3. After brief coomassie blue staining of the gels, lanes were cut into low, medium and high molecular weight segments, excluding the globin band, and processed for in-gel trypsin digestion ^27^. The extracted tryptic peptides were dissolved in Solvent A (95/5% water/acetonitrile containing 0.1% formic acid); and 1 μl was injected for analysis by LC-MS/MS. The nanoLC-MS consisted of an UltiMate 3000 Nano LC System and an LTQ-Orbitrap Elite mass spectrometer (Thermo Fisher, San Jose, CA). Reversed-phase liquid chromatography was performed using a homemade 33 cm× 75 μm ID column packed with XBridgeTM BEH C18 media (2.5 μm, 130 Å). The flow rate was maintained at 200 nl/min. Solvents A and B (95/5% acetonitrile/water containing 0.1% formic acid) were used to establish the 160 min gradient elution timetable: isocratic at 5% B for 30 min, 5-55% B over 70 min, followed by 55-99% B in 5 min where it was maintained for 10 min, and finally returned to 5% B over 5 min for a 40 min re-equilibration time. The LTQ-Orbitrap Elite mass spectrometer instrument was operated in positive mode with a 2.6 kV applied spray voltage. The temperature of the ion transfer capillary was 300 °C. One microscan was set for each MS and MS/MS scan. A full scan MS acquired in the range 300 ≤ m/z ≤ 2000 was followed by 10 data dependent MS/MS events on the 10 most intense ions. The mass resolution was set at 60000 for full MS. The dynamic exclusion function was set as follows: repeat count, 1; repeat duration, 30 s; exclusion duration, 30s. HCD was performed using normalized collision energy of 35% and the activation time was set as 0.1 ms. The tryptic peptides were separated by nanoflow HPLC and analyzed by an ESI-LTQ-orbitrap mass spectrometer. The sequencing data was searched against the NCBI *I. scapularis* protein database (accessed on June 30, 2020) using Mascot (Matrix Science) for protein identification and quantification. The search parameters were: peptide mass tolerance ± 25 ppm, fragment mass tolerance ± 50 mmu, maximum of 3 missed cleavages.

Individual gel bands of purified recombinant TULIP2 were excised with a blade and subjected to in-gel trypsin digestion. TULIP2-His was analyzed by liquid chromatography-tandem mass spectrometry (LC-MS/MS) as described above. MBP-TULIP2 was analyzed by capillary HPLC-electrospray ionization tandem mass spectrometry on a Thermo Scientific Orbitrap Fusion Lumos mass spectrometer. On-line HPLC separation was accomplished with an RSLC NANO HPLC system (Thermo Scientific/Dionex) interfaced with a Nanospray Flex ion source (Thermo Scientific): column, PepSep (Bruker; ReproSil C18, 15 cm x 150 µm, 1.9 µm beads); mobile phase A, 0.5 % acetic acid (HAc) / 0.005 % trifluoroacetic acid (TFA) in water; mobile phase B, 90 % acetonitrile / 0.5 % HAc / 0.005 % TFA / 9.5 % water; gradient 3 to 42% B in 60 min; flow rate, 800 nl/min. Precursor ions were acquired in the orbitrap in centroid mode at 120,000 resolution (*m/z* 200); data-dependent higher-energy collisional dissociation (HCD) spectra were acquired at the same time in the linear trap using the “rapid” speed option (30% normalized collision energy). Other MS scan parameters included: mass window for precursor ion selection, 0.7; charge states, 2 – 5; dynamic exclusion, 15 sec (± 10 ppm); intensity to trigger MS2, 50,000. Mascot (v3.0.0; Matrix Science, London UK) was used to search the spectra against UniProt_E_coli_ref 83333_20220830 (4,402 sequences; 1,354,438 residues) and some custom databases. The following search parameters were used: peptide tolerance 20 ppm, fragment tolerance 0.8 Da; variable modifications including cysteine carbamidomethylation, methionine oxidation and deamidation of glutamine and asparagine; two missed tryptic cleavages allowed.

Protein quantification was calculated with intensity based absolute quantification (iBAQ) ^28^, implemented using custom software and Excel spreadsheets. For each observed protein detected, iBAQ is the total intensity of the observed peptides divided by the number of theoretically observable peptides for that protein. Peptide intensities were obtained from the raw mass spectrometer output using Mascot (Matrix Science) software to match spectrometer scans with identified peptides. Only peptides with Mascot scores having identification probabilities of p>0.95 were included in the analysis. The intensities were combined for each identified protein in the three gel slices in each sample. iBAQ values were normalized for each protein in separate biological replicates using the total intensity for all peptides in that replicate. Statistical comparisons of iBAQ were done using JMP Pro 17 software (Cary, NC).

### 2.11 Lipid analysis

High-resolution mass spectra were collected on a maXis plus quadrupole-time of flight mass spectrometer equipped with an electrospray ionization source (Bruker Daltonics, Billerica, MA) and operated in negative ionization mode. Lipid extracts from whole cells, solubilized in chloroform (see section 2.6), were introduced into the ESI source via syringe pump at a constant flow rate of 3 μL/min. Important source parameters are summarized as follows: capillary voltage, –3500 V; nebulizer gas pressure, 0.4 bar; dry gas flow rate, 4.0 L/min; source temperature, 200 °C. Mass spectra were averages of one minute of scans collected at a rate of 1 scan per second in the range 50 ≤ m/z ≤ 1500. Compass Data Analysis software version 4.3 (Bruker Daltonics) was used to process all mass spectra. Peaks were identified and annotated in the software mMass ^29,30^.

### 2.12 Computational analysis

The *I. scapularis* salivary TULIPs, initially found by proteomic analysis, are annotated by the National Center for Biotechnology Information (NCBI) as “uncharacterized” proteins (https://www.ncbi.nlm.nih.gov/). We identified them as members of the TULIP superfamily using SwissModel homology modeling (https://swissmodel.expasy.org/). Paralogs were found using BLAST software (https://blast.ncbi.nlm.nih.gov/Blast.cgi). After trimming predicted signal sequences from the TULIP N-termini with SignalP software (https://services.healthtech.dtu.dk/services/SignalP-6.0/), we constructed an evolutionary tree of the TULIP paralogs using raxmlGUI 2.0 (https://antonellilab.github.io/raxmlGUI/). Molecular models of TULIPs were generated with the online AlphaFold3 server (https://alphafoldserver.com) and RoseTTAFold (https://robetta.bakerlab.org/), using the signal-trimmed NCBI amino acid sequences. Internal pockets in the structures were predicted using CASTp software (http://sts.bioe.uic.edu/castp/calculation.html). Analysis of the molecular models for predicted membrane-insertion sites was done using DREAMM software (https://dreamm.ni4os.eu/).

Ligand docking was calculated with AutoDock Vina software (vina_1.2.7_mac_aarch64) ^31,32^. Ligand structures were obtained from PubChem in sdf format and converted to PDB format using PyMOL (https://pymol.org). The TULIP molecular models and ligand structures were converted to pdbqt format using Open Babel (https://openbabel.github.io). The docking grid box was drawn to include the entire TULIP molecule, using Webina 1.0.5 (https://durrantlab.pitt.edu/webina/). Vina ran with a completeness parameter of 32 for lipids containing cholesterol and 16 for lipids containing two acyl chains. Ligand docking was also done with Boltz2 (https://labs.rowansci.com/) and Alphafold3 running on a local high-performance computer cluster (https://github.com/google-deepmind/alphafold3).

### 2.13 Molecular dynamics

We employed VMD ^33^ to place the protein in a 100 Å × 100 Å× 100 Å box of water (TIP3P). The system was then neutralized and salinated with Na⁺ and Cl^−^ ions to a salt concentration of 150 mM. We employed NAMD 2.13 (for initial equilibration) and 3.0 (for production runs) ^34,35^ as the MD engines. We used CHARMM36 parameters ^36–38^ for inter-and intra-molecular interactions. After initial equilibration of the system, we ran unbiased MD for 10 ns with constant pressure at 1.0 bar (Nose-Hoover barostat) and constant temperature at 298 K (Langevin thermostat). Subsequently, we ran unbiased MD for 400 ns with constant pressure at 1.0 bar and constant temperature at 380 K. After that, we ran unbiased MD for 220 ns with constant pressure at 1.0 bar and constant temperature at 450 K. In all the simulation runs, the Langevin damping coefficient was chosen to be 1/ps. The periodic boundary conditions were applied to all three dimensions. The particle mesh Ewald (PME) was used for the long-range electrostatic interactions (grid level: 128×128×128). The time step was 2.0 fs. The cut-off for long-range interactions was set to 10 Å with a switching distance of 9 Å.

## 3. Results

### 3.1 *I. scapularis* salivary proteome

The salivary proteome (sialome) from pilocarpine-induced saliva was measured on nine *I. scapularis* nymphs, five that had fed as larvae on uninfected mice and four fed as larvae on *B. burgdorferi*-infected mice. Six members of the TULIP family were identified in saliva (Figure 1). We detected 252 proteins that were present in four out of five uninfected or three out of four infected nymphs. They are displayed as a “volcano” plot in Figure 2, with the normalized difference in protein amounts (measured as intensity based absolute quantification, iBAQ) between saliva of tick nymphs that had fed as larvae on mice that were uninfected U or *B. burgdorferi*-infected mice I, (U-I)/(U+I), averaged for the ticks in each group and plotted against the logarithm of the statistical significance of the difference. Twelve proteins showed statistically significant differences (P<0.05) in a t-test: four that significantly increase in the presence of *B. burgdorferi* and eight that significantly decrease (Table 1). Of the increasing proteins, two are proteases and two are protease inhibitors. Of the decreasing proteins, four are known in other organisms to have activities involving lipids. One of these is a TULIP, which we refer to as TULIP2 (NCBI accession number XP_029833768.1; currently called XP_040079658.2; see Supporting Information Figure S1 for a discussion of observed polymorphisms). TULIP2 showed a significantly lower abundance in nymphs that had fed as larvae on *B. burgdorferi*-infected mice.

**Figure 1.**
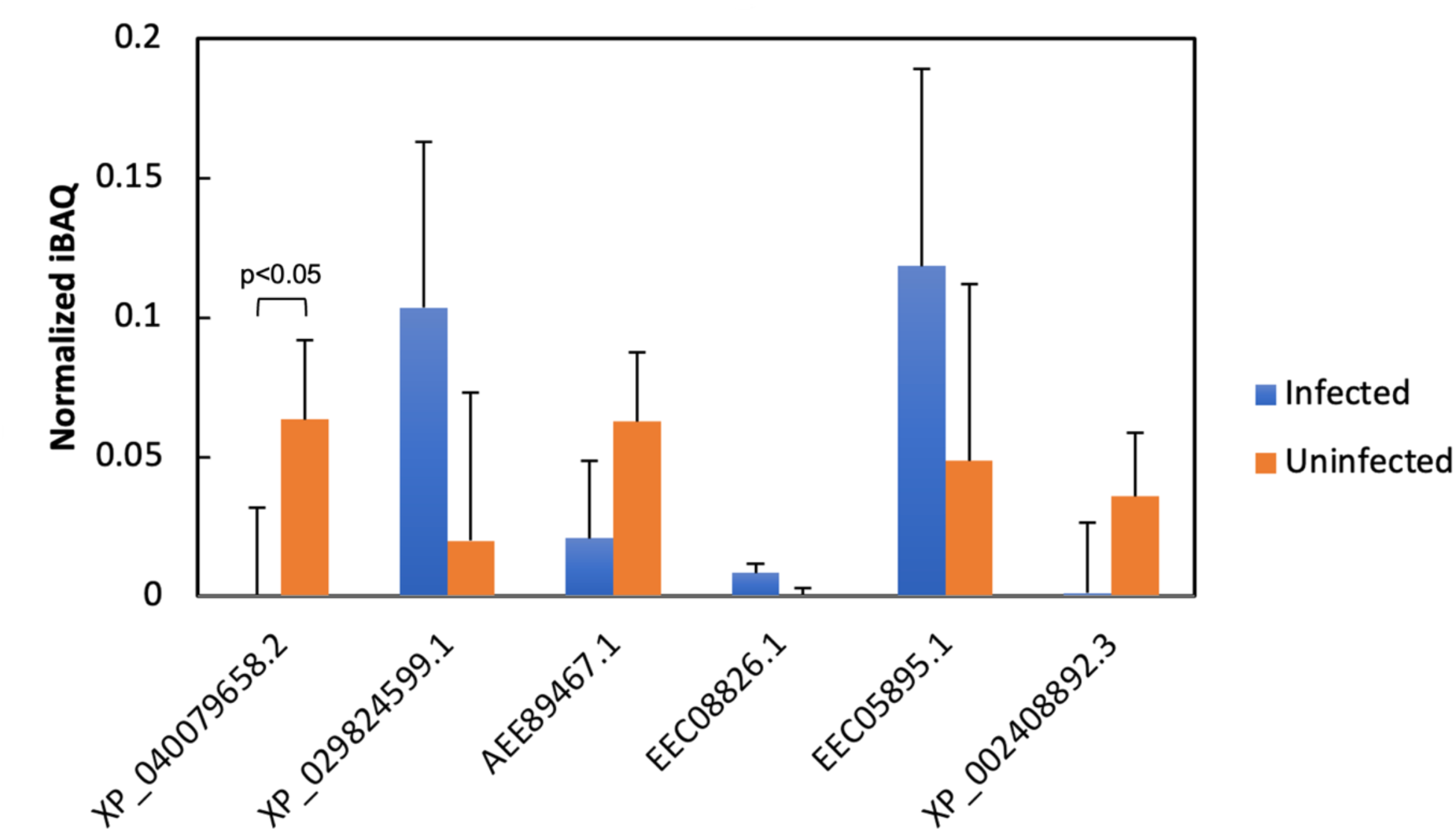
Tubular lipid-binding proteins (TULIPs) in *I. scapularis* saliva. TULIPs identified in saliva of *B. burgdorferi* -infected (blue) and non-infected (orange) *I. scapularis* nymphs. Protein levels calculated by intensity-based absolute quantification (iBAQ), normalized to total protein intensity in each sample (see Methods). Bars show the mean of 5 uninfected or 4 infected nymphs. Error bars indicate the standard deviation. Difference between protein levels in uninfected and infected nymphs was statistically significant at the p<0.05 level for XP_040079658.2, TULIP2.

**Figure 2.**
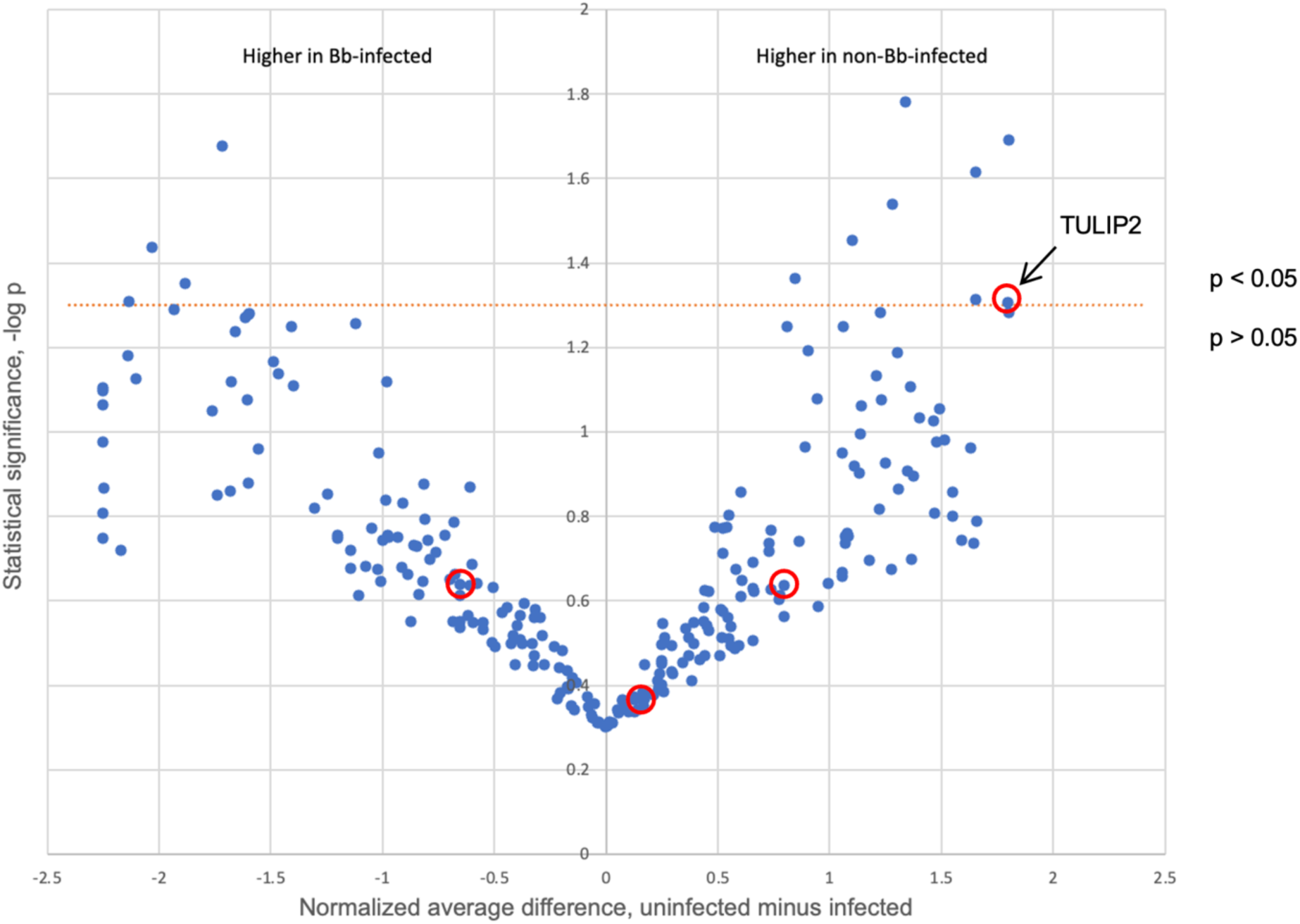
Volcano plot of *I. scapularis* sialome, comparing expression levels in saliva of nymphs fed on uninfected vs. *B. burgdorferi* -infected mice. Protein levels calculated by intensity-based absolute quantification (iBAQ). Statistical significance calculated using t-test. See text for details. p<0.05 level indicated by dotted red line. Circles: four single-domain tubular lipid-binding proteins (TULIPs) that were highly expressed in nymph saliva. Only TULIP2 (arrow) had statistically significant differences between *B. burgdorferi*-infected and uninfected nymphs.

**Table 1.** Salivary proteins that are significantly expressed at higher levels in *I. scapularis* nymphs feeding on uninfected mice compared with *B. burgdorferi*-infected mice.

| Accession ID | -log p<sup>a</sup> | Average intensity difference<sup>b</sup> | Protein |
| --- | --- | --- | --- |
| XP_029822532.1 | 1.78251606 | 1.34 | vitellogenin-5 |
| EEC16393.1 | 1.69036983 | 1.80 | annexin V |
| XP_029838979.1 | 1.67778071 | -1.72 | cathepsin L |
| XP_002406560.2 | 1.61439373 | 1.65 | heat shock protein |
| XP_029838392.1 | 1.53910216 | 1.28 | saposin-like |
| XP_029843550.1 | 1.45345734 | 1.10 | chitinase 10 |
| EEC18341.1 | 1.43651891 | -2.03 | trypsin inhibitor |
| EEC14101.1 | 1.36251027 | 0.84 | aldolase |
| EEC14237.1 | 1.35066514 | -1.88 | serpin |
| XP_029825410.1 | 1.31336373 | 1.65 | phosphoserine aminotransferase |
| XP_029823653.1 | 1.30891851 | -2.13 | chymotrypsin-elastase inhibitor |
| XP_029833768.1 | 1.30627305 | 1.80 | TULIP2 |

Footnotes: a) Probability p that protein level is significantly different in saliva of nymphs feeding on *B. burgdorferi*-infected mice compared with saliva of ticks feeding on uninfected mice, based on t-test. b) Expression difference using intensity-based quantification (iBAQ; see Methods): uninfected minus infected.

Numerous TULIP paralogs previously have been identified in insect genomes ^39^. To find whether ticks also have a large paralogous group of TULIP genes, we used BLAST searches to identify other *I. scapularis* TULIPs. We found 33 TULIP paralogs (Figure 3) that differed by more than six substitutions and also were likely to be secreted (i.e. predicted to have N-terminal signal sequences). The list does not include two multi-domain proteins with C-terminal TULIPs: one having an N-terminal sulfotransferase sequence (EEC08826.1) that we detected in saliva (Figure 1), and another with N-terminal immunoglobulin domains (XP_040079652.1) annotated by NCBI as neuroglian. A tree of *I. scapularis* TULIP paralogs is shown in Figure 3. The TULIPs found in the sialome (this work and Kim et al. ^40^) are distributed primarily in branches near TULIP2, but several occur in more distant branches.

**Figure 3.**
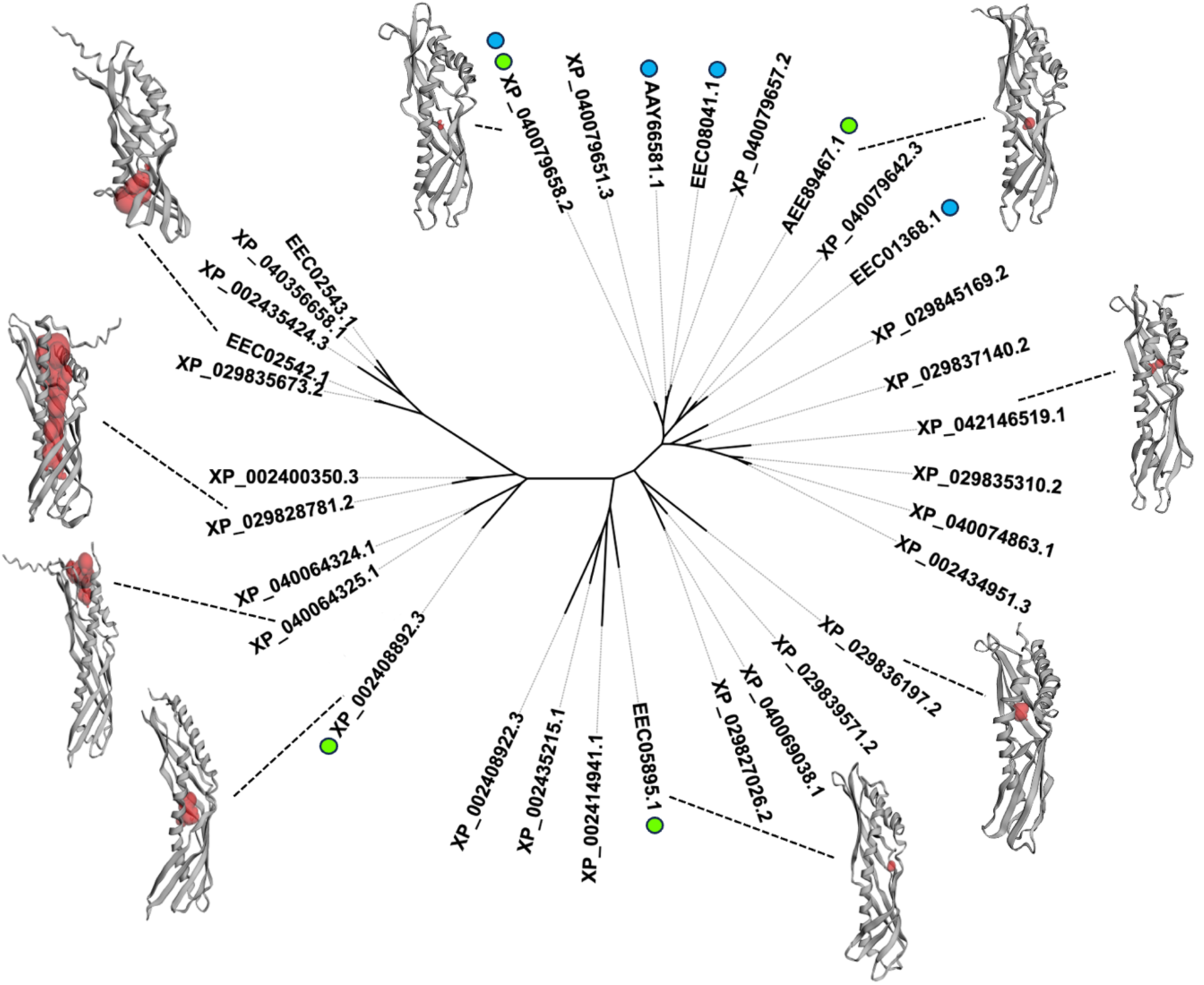
Tree of *I. scapularis* TULIP paralogs. Unrooted tree of 32 TULIPs from *I. scapularis* displayed using iTOL (https://itol.embl.de/). Branches are identified by protein sequence accession numbers. Branch lengths are proportional to amino acid sequence differences. Circles next to accession numbers mark TULIPs identified in the *I. scapularis* sialome: green, this work; blue, Kim et al., 2021. Protein backbone cartoons, with dashed lines to the accession numbers, are from AlphaFold3 structure predictions. The largest interior cavity is shown in red, as calculated by CASTp. For clarity, XP_029824599.1 was omitted from the tree because of its proximity to XP_040079658.2.

### 3.2 Expression and purification of TULIP2-His

Considering the involvement of some TULIP family members in immunity, we sought to measure the interaction of TULIP2 with *B. burgdorferi* lipids. TULIP2, with a C-terminal 6x-His tag (TULIP2-His), was expressed primarily in *E. coli* inclusion bodies, which were separated from the cell lysate by centrifugation and dissolved in 5 M guanidinium hydrochloride (Gdn). Further purification on a nickel affinity column (Figure S2), showed the main product at the expected size of recombinant TULIP2-His (25 kDa), as well as higher molecular weight bands, confirmed by mass spectrometry to be TULIP2-His and TULIP2-His multimers.

TULIP2-His precipitated after dialysis against phosphate-buffered saline (PBS) or PBS containing decreasing concentrations of Gdn to gradually lower the Gdn concentration in the protein sample. Centrifugal ultrafiltration filters (10 kDa molecular weight cutoff) were also tried for exchanging Gdn, imidazole and excess NaCl, but TULIP2-His aggregated on the filter membrane.

Various additives and conditions were considered and tested to aid TULIP2-His refolding, including low and high pH (Figure S3), detergent bicelles, and disulfide exchange (Figure 4). Solubilization of TULIP2-His was not observed in any of these conditions. However, disulfide exchange appears to explain part of the aggregation mechanism. TULIP2 contains two cysteine residues in its amino acid sequence, modeled by AlphaFold3 as a disulfide (see section 3.6). A cystine-cysteine redox refolding system was employed in an attempt to break and reform disulfide bonds and promote the native folded conformation of TULIP2^41^. The redox equilibrium was set by 1 mM cystine and 12 mM or 17.9 mM cysteine in 100 mM Tris, 200 mM NaCl, pH 8.0, with or without 10% glycerol. TULIP2-His at 59 µM was heated with 0.5 mM DTT to 30 °C for 30 min and then cooled to room temperature and left for 10 min. The protein sample and the cystine-cysteine redox refolding buffer with or without glycerol was mixed at a 1:1 ratio, and allowed one hour to equilibrate at room temperature. Subsequently, the four samples were dialyzed against Tris buffer containing the cystine-cysteine pair at a lower concentration. After the first two hours of dialysis, protein aggregation was obvious (Figure 4). When DTT was omitted from the SDS-PAGE loading buffer, a ladder of higher molecular-weight bands was observed. This suggests that intermolecular disulfide bonding contributes to the aggregation of TULIP2-His.

**Figure 4.**
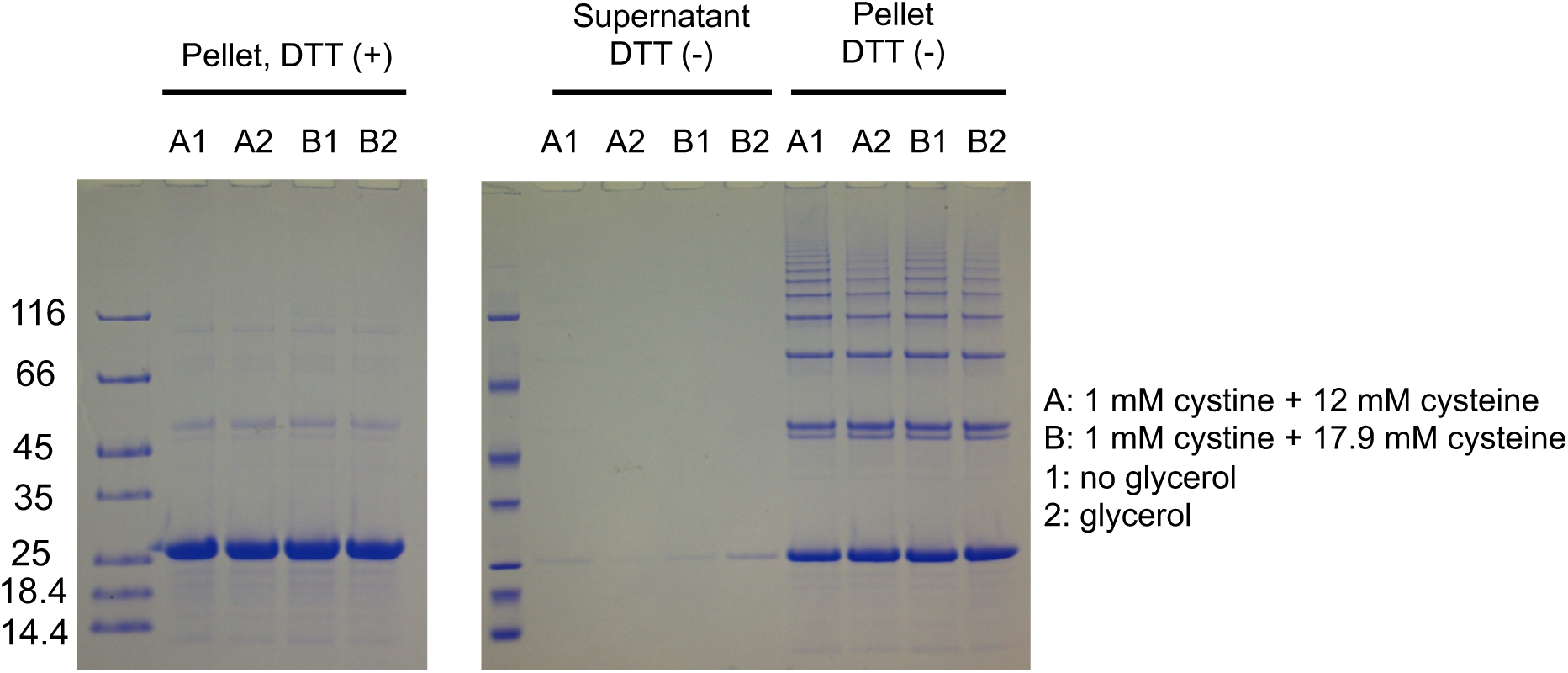
Cystine-cysteine redox refolding of TULIP2. TULIP2 was refolded by dialysis in the indicated cystine/cysteine concentrations, in the presence or absence of 10 % glycerol. Gels were run with or without DTT for the pellet samples

Mild detergents, including n-dodecyl β-ᴅ-maltoside (DDM) and octyl glucoside (OG) have been used for refolding unfolded proteins or proteins expressed in inclusion bodies ^42,43^. To test whether TULIP2 can be refolded with chemical additives, TULIP2-His in 5 M Gdn was rapidly diluted 5-fold by adding the protein sample into diluents to obtain a final concentration of 1 M Gdn and additives as follows: 2 % CHAPS, 1 % CHAPS, 1.5 % OG and 2 % DDM. The diluted protein samples were allowed to equilibrate at room temperature for 2.5 hours. Absorbance at 330 nm was measured to determine the turbidity of the protein samples after rapid dilution. The samples were centrifuged, and the absorbance of the supernatant at 280 nm was measured to determine the soluble protein remaining in solution. The results (Table S1, Figure S4) show that DDM was the most effective in mitigating protein aggregation. However, it was not possible to remove DDM and Gdn and still maintain TULIP2 solubility.

### 3.3 Expression and purification of MBP-TULIP2

To address the solubility and refolding issue, another vector was constructed to express TULIP2 with an N-terminal maltose-binding protein (MBP) tag ^44^. The MBP tag made it possible to recover soluble recombinant protein from the supernatant of cell lysis materials without additives. The fractions of the MBP-trap affinity column purification were examined by SDS-PAGE (Figure S5), and confirmed by western blot using two different primary antibodies). The two primary antibodies both revealed a band at the expected size 65 kDa (Figure S6). The successful expression of MBP-TULIP2 was also confirmed by mass spectrometry.

Next, attempts were made to cleave the MBP tag with TEV protease, which has a His tag for Ni affinity removal. Regardless of whether MBP or TEV protease was removed first, the tag-free TULIP2 protein disappeared after the first removal step. Also, it was found that in the maltose-affinity purification, elution by maltose was incomplete, even if the concentration was raised to 20 mM. The incomplete elution by maltose observed during maltose-affinity purification of MBP-TULIP2 from bacterial lysate, and the fact that TULIP2 adsorbs to the 6 % crosslinked agarose-based MBPTrap HP column and Ni-NTA resin suggest that TULIP2 may have an affinity for galactosides. To test this hypothesis, another MBP-TULIP2 purification by MBPTrap HP column was performed. After the standard maltose elution, 300 mM lactose in addition to 10 mM maltose was applied to the column. SDS-PAGE analysis showed more MBP-TULIP2 was eluted after adding lactose, although the elution was still incomplete (Figure S7).

### 3.4 TULIP2 selectively recognizes *B. burgdorferi* lipids

Because many TULIPs bind lipids, and the level of TULIP2 in saliva decreases in *B. burgdorferi*-infected nymphs, we asked whether TULIP2 interacts with *B. burgdorferi* lipids. A protein-lipid overlay assay was carried out ^45,46^, detecting TULIP2 with mouse anti-TULIP2 antiserum, followed by horseradish peroxidase-conjugated goat anti-mouse IgG secondary antibody and chemiluminescence. A chloroform-methanol extract of lipids from *B. burgdorferi* was used in the overlay assay. The composition of the extract was examined by mass spectrometry. It has been reported that *B. burgdorferi* lipid comprises phosphatidylcholine (PC), phosphatidylglycerol (PG), mono-α-galactosyl-diacylglycerol (MGalD), cholesteryl-β-D-galactopyranoside (CGal), cholesteryl 6-O-acyl-β-D-galactopyranoside (ACGal) and free cholesterol and cholesteryl esters ^23–25^. In our *B. burgdorferi* lipid extract we identified by mass spectrometry PC (acyl chains totaling 34 carbons) and CGal (Figures S8-S11).

The *B. burgdorferi* lipid extract showed a TULIP2 binding signal (Figure 5, Figure S12) whereas cholesteryl stearate and 1-palmitoyl, 2-oleoyl phosphatidylcholine (POPC) did not. PC constitutes about 10% of *B. burgdorferi* lipids ^23^. We cleaved the MBP tag from MBP-TULIP2 using TEV protease and showed that the tag did not interfere with lipid binding (Figure S12). A blank blot without TULIP2 dots and not incubated with TULIP2 showed no chemiluminescence signal, indicating that the antibodies themselves do not bind to the lipids. These blots demonstrate that TULIP2 recognizes one or more specific lipid components of *B. burgdorferi*. We tested two additional lipids: 1,2-dipalmitoyl-sn-glycerol and D-galactosyl-α-1,1′ N-palmitoyl-D-erythro-sphingosine (C16 galactosyl(⍺) ceramide). A previous report showed that 1-palmitoyl-2-oleoyl diacylglycerol elicits IMD responses in ticks ^9^; and the α-galactosyl group is found in the *B. burgdorferi* lipid mono-α-galactosyl-diacylglycerol ^25^. Only the extracted *B. burgdorferi* lipids showed a strong TULIP2 binding signal (Figure 5). The effect of the presence of the MBP tag was tested by comparing MBP-TULIP2 and TEV protease-cleaved MBP-TULIP2.

**Figure 5.**
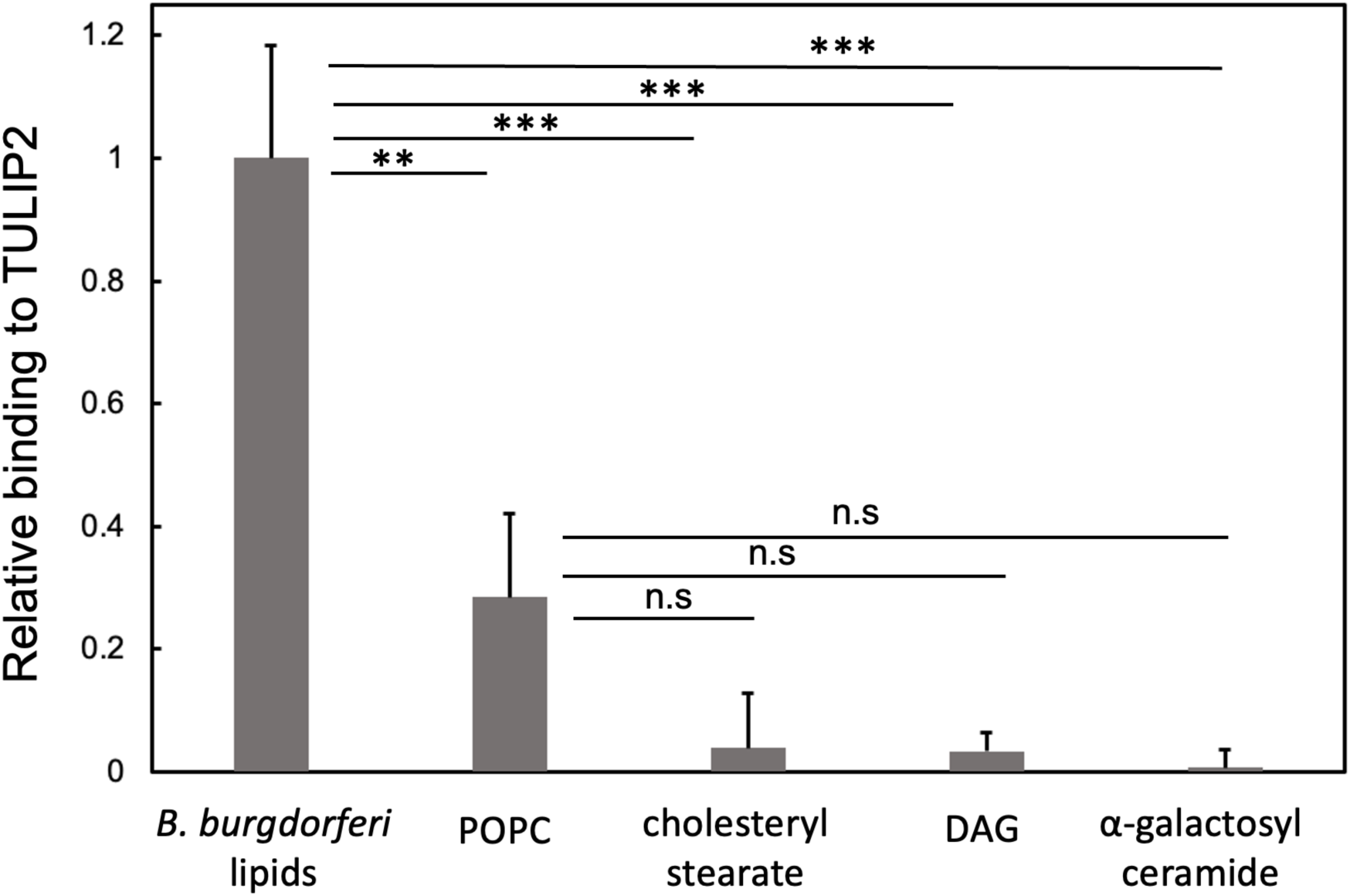
Binding of TULIP2 to *B. burdorferi* lipid extract and control lipids. Extracted *B. burgdorferi* lipids (∼0.7-0.8 nmol total) were spotted on nitrocellulose films, along with spots of palmitoyl oleoyl phosphatidylcholine (POPC) (39 nmol), cholesteryl stearate (47 nmol), 1,2-dipalmitoyl-sn-glycerol (DAG) (80 nmol), and ⍺-galactosyl ceramide (21 nmol). After blocking with bovine serum albumin, films were equilibrated with TULIP2, washed, developed with mouse anti-TULIP2 antibody and horseradish peroxidase-linked anti-mouse IgG, and imaged by chemiluminescence. Spot intensities, quantified by ImageJ, were normalized to *B. burgdorferi* lipids. Statistical significance (t-test): **, P < 0.005; ***, P< 0.0005; n.s., not significant

### 3.5 Molecular modeling

AlphaFold3 models were built for the TULIPs in Figure 3, using the AlphaFold Server. The best-scoring models were uploaded to the CASTp server to predict internal cavities. None of the salivary-expressed TULIPs were predicted to have large interior cavities suitable for lipid binding (Figure 3). For example, with the AlphaFold3 model of TULIP2 (Figure 6A and B), AutoDock predicts cholesteryl-β-D-galacto-pyranoside binds only to surface sites, with free energies less than 7 kcal/mol, implying a dissociation constant of greater than 8 μM. Although AI diffusion methods can use receptor flexibility to model ligand-protein complexes ^47–49^, we obtained only predictions of surface binding sites from the full AlphaFold3 software run on a local server, and Boltz2 run on a web server. The apparent failure to detect ligand-binding pockets may reflect the technical limitations of using AI-derived models for ligand docking predictions. The opening of an interior ligand-binding site typically involves protein conformational changes, which AI modeling does not yet handle well ^50,51^. As an alternative, we used molecular dynamics to explore TULIP2 conformations near the AlphaFold3-predicted lowest energy structure, by simulating thermal unfolding ^52^.

**Figure 6.**
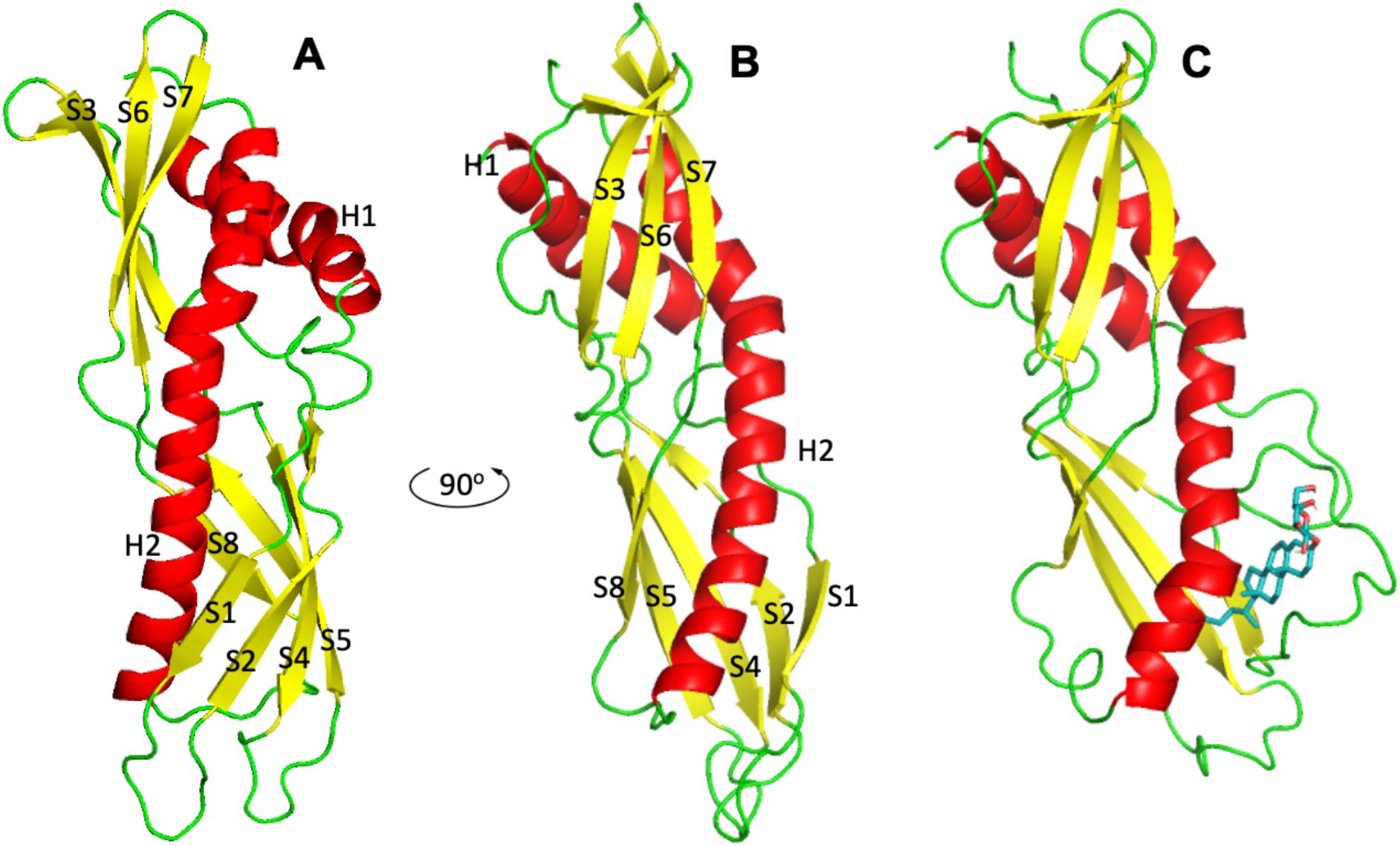
Models of TULIP2. A) AlphaFold3 model of TULIP2, oriented to show the S1-S2 and S4-S5 loops. B) Rotation of A 90° to the right, showing the seam between helix H2 and β strand S1. C) Model from A and B heated to 450 K for 43.8 nsec in a molecular dynamics simulation. The H2-S1 seam opens a hydrophobic pocket. AutoDock Vina predicts this pocket has a high affinity for cholesteryl-β-D-galactopyranoside (shown in cyan stick representation). Protein color coding: α-helix, red; β strand, yellow; loops and undefined secondary structure, green.

The AlphaFold3 model for TULIP2 was computationally heated to 450 K, and molecular dynamics were followed for 100 nsec (Figure 7). After about 30 nsec, separation began along the seam between α helix H2 (positions 160-180) and β strand S1 (positions 25-40), opening a hydrophobic pocket (Figure 6). By contrast, over the same time period, very little change occurs in the distance between the N-terminal α-helix H1 (positions 10-25) and the β-sheet formed by S6 and S7 (positions 120-140) adjacent to the last part of the C-terminal α-helix H2 (Figure 6). The thermally-opened seam appears to be the least stable feature of the folded TULIP2 structure. If TULIIP2 can form an internal binding pocket similar to known TULIP structures, the exposed pocket may be similar to a TULIP2 conformational intermediate in lipid binding. We used AutoDock to model binding of cholesteryl-β-D-galactopyranoside to the TULUP2 conformation that occurred at 43.8 nsec. The results show strong binding of cholesteryl-β-D-galactopyranoside (Cgal) to the exposed pocket, with an apparent dissociation constant of 0.44 μM, an interaction about twenty-fold stronger than for the predicted binding sites on the surface of the 25°C structure (Table 3). Of the *B. burgdorferi* lipids examined by computational docking, Cgal had the strongest predicted binding to the 43.8 nsec heated TULIP2 structure. Phosphatidylcholine (PC) was nearly 7-times weaker, which is consistent with the experimental dot blot results (Figure 5). We also tested docking of phosphatidylserine and phosphatidylethanolamine (see Discussion), and these lipids showed weaker binding than PC.

**Figure 7.**
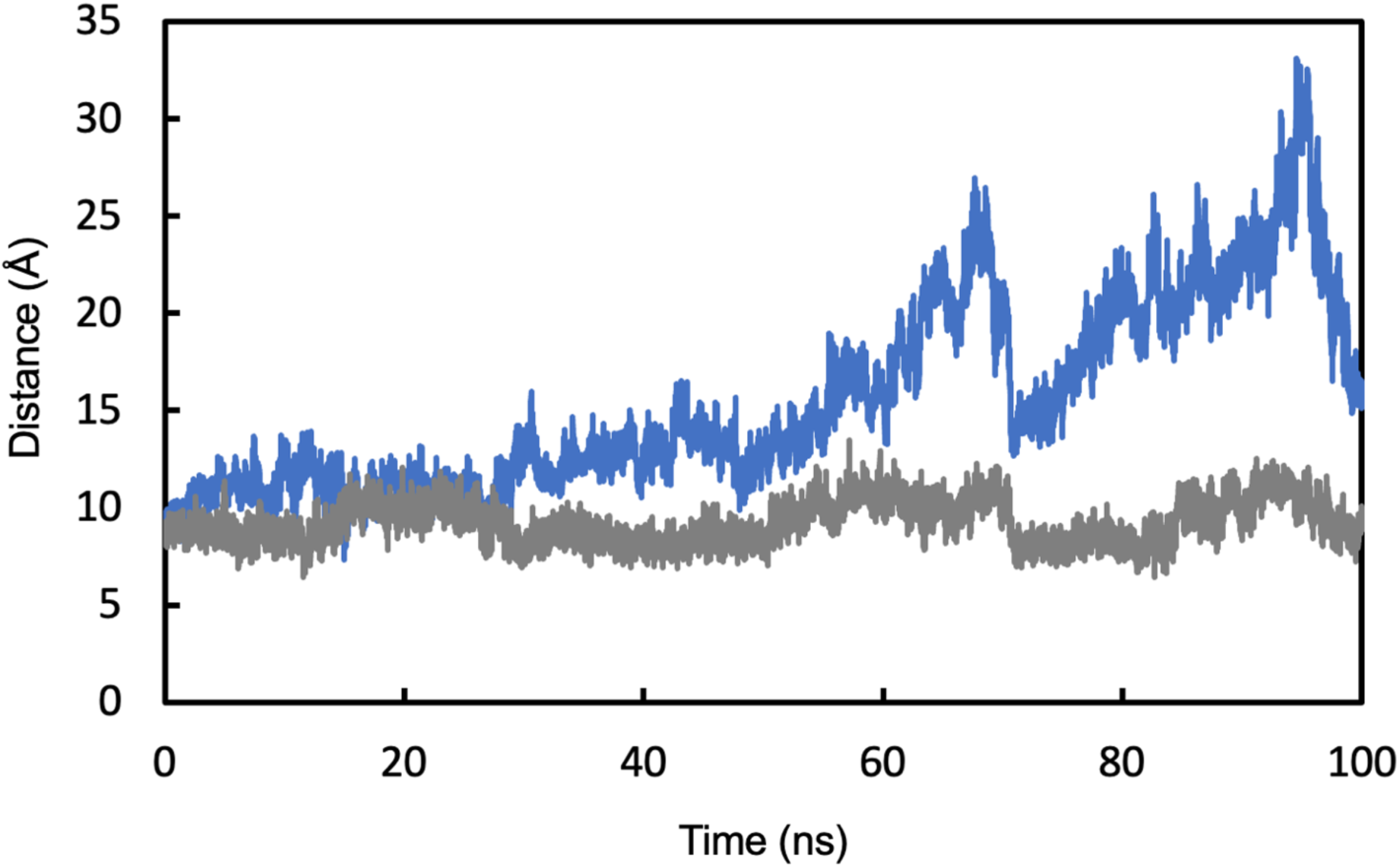
Molecular dynamics simulation of rapid heating of TULIP2. The AlphaFold3 model of TULIP2 was heated to 450K. After 30 nsec, the centroid of amino acids 160-180 begins to separate from the centroid of amino acids 25-40 (blue line), but little change was observed over the same time period for the distance between the centroids of amino acids 10-25 and 120-140 (gray line).

**Table 2.**
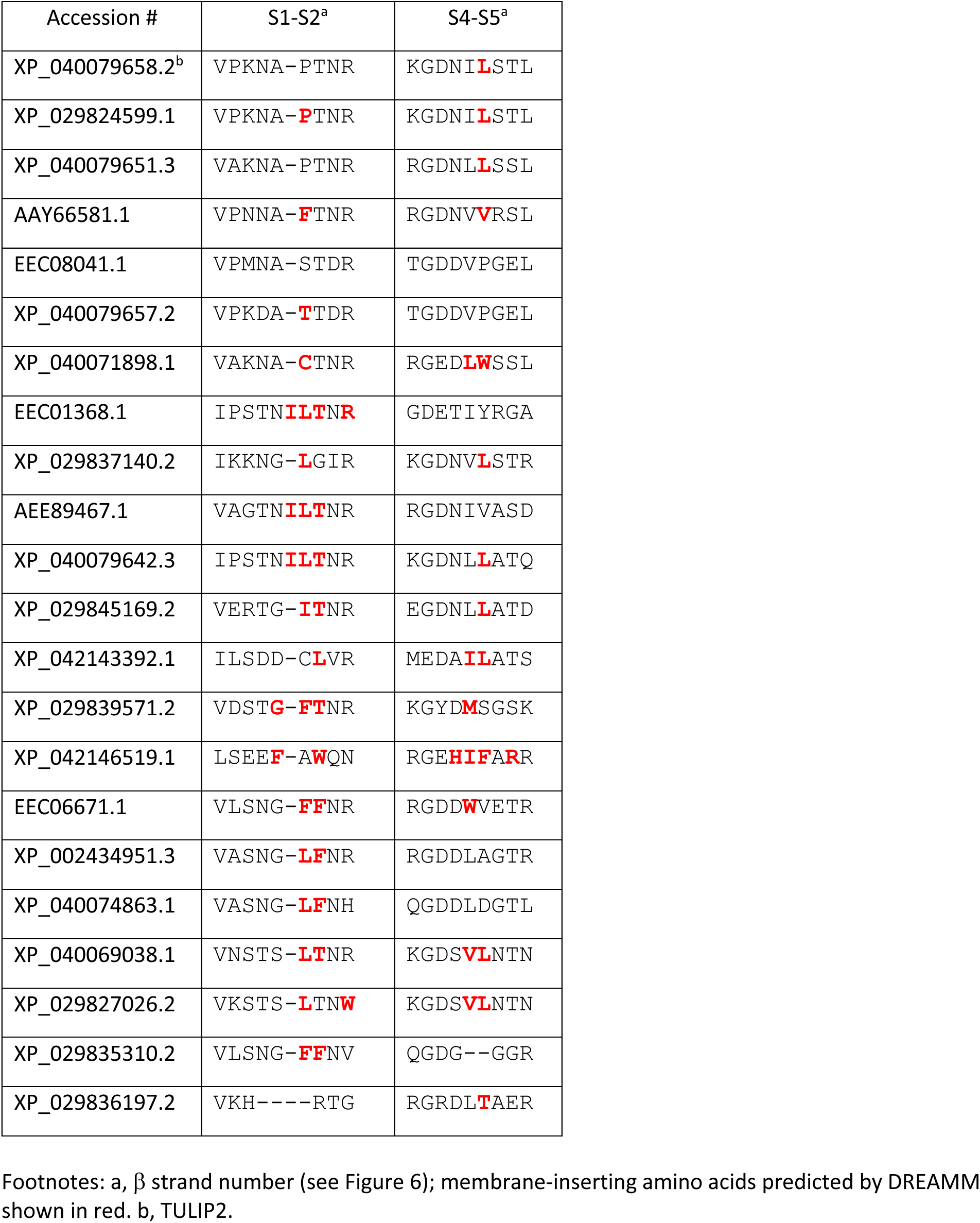
Predicted membrane-inserting sequences in *I. scapularis* TULIPs.

**Table 3.** Computational ligand docking of *B. burgdorferi* membrane lipids to TULIP2 molecular models.

| Model<sup>a</sup> | Cgal<sup>b</sup> |  | ACGal<sup>c</sup> |  | PC<sup>d</sup> |  | MGalD<sup>e</sup> |  |
| --- | --- | --- | --- | --- | --- | --- | --- | --- |
| | $\Delta G^{\circ}$<sup>f</sup> | $K_d$<sup>g</sup> | $\Delta G^{\circ}$ | $K_d$ | $\Delta G^{\circ}$ | $K_d$ | $\Delta G^{\circ}$ | $K_d$ |
| AF3 | -6.91 | 8.2 | -4.64 | 384 | -3.31 | 3660 | -5.52 | 87 |
| MD | -8.63 | 0.44 | -7.37 | 3.8 | -7.51 | 3.0 | -7.76 | 1.9 |

Footnotes: a, AF3 model of TULIP2 from AlphaFold3 Server; MD model of TULIP2 AF3 model after heating to 450 K for 43.8 nsec. b, cholesteryl-β-D-galactopyranoside. c, cholesteryl 6-O-stearoyl- β-D-galactopyranoside. d, 1-palmitoyl-2-oleoyl-glycero-3-phosphocholine. e, mono-α-galactosyl-diacylglycerol. f, predicted binding standard free energy change, Kcal/mol. g, dissociation constant calculated at 25°C from $\Delta G^{\circ}$ , μM.

Although structures such as the ones occurring after >30 nsec of heating appear to have strong interactions with *B. burgdorferi* lipids, it is not clear how a conformational change to this structure would occur under biologically relevant conditions. Molecular dynamics trajectories at 298 K do not detect this conformational change over a 100 nsec time scale. However, at ordinary temperatures, we do observe a significant motion of the loops connecting β strands S1 and S2, and β strands S4 and S5. A possible trigger for opening a TULIP2 lipid-binding pocket could be interaction with lipid bilayers, interactions known in other TULIPs ^53^. We used DREAMM software to predict membrane-binding loops in TULIP2. The results (Table 2) show that the S4-S5 loop of TULIP2 is likely to be membrane-inserting. Nearly all the *I. scapularis* TULIP paralogs were predicted by DREAMM to bind membranes via these loops (Table 2). Additional membrane binding sites were predicted for some of the *I. scapularis* TULIP paralogs at the S3 and S6-S7 loops, which are at the opposite end of the TULIP molecule from the S1-S2 and S4-S5 loops (Figure 6), suggesting that some *I. scapularis* TULIPs interact with two different membranes.

## 4. Discussion

### 4.1 *I. scapularis* TULIP2 interaction with *B. burgdorferi* lipids

This work clearly shows that the *I. scapularis* salivary protein TULIP2 binds to *B. burgdorferi* lipids (Figure 5). This suggests the possibility that TULIP2 recognizes *B. burgdorferi* lipids as part of *I. scapularis* immune pathways.

We have not definitively identified the lipid or lipids that bind to TULIP2. However, our results suggest it could be cholesteryl-β-D-galactopyranoside. We did not detect any TULIP2 binding to cholesteryl stearate or to ⍺-galactosyl C16 ceramide (Figure 5). However, during purification of TULIP2, we found evidence that TULIP2 binds to agarose (Figure 4), which is a polymer containing β-galactosyl linkages. TULIP2 is released from agarose by excess lactose, a β-galactoside. The analysis of our *B. burgdorferi* lipid extract showed significant amounts of cholesteryl-β-D-galactopyranoside, which is likely to be the major galactoside lipid in our extract, as we did not detect mono-α-galactosyl-diacylglycerol or cholesteryl 6-O-acyl-β-D-galactopyranoside (Figures S8 and S11).

Selective binding by TULIP2 to cholesteryl-β-D-galactopyranoside is supported by computational analysis. The structure prediction models for TULIP2 and the other salivary TULIPs (Figure 3) do not have internal binding pockets, and only weak binding to lipids was predicted. Ligand binding to proteins often involves protein conformational changes, which AI-based algorithms are not currently able to predict reliably. To overcome this, we used molecular dynamics simulation to thermally drive the AlphaFold3-predicted structure of TUILP2 into slightly less stable conformational states (Figure 7). After about 30 nsec of heating, we observed a structure that had a hydrophobic pocket (Figure 6C), which may resemble a conformational transition state involved in ligand binding. This conformation showed strong and selective binding to cholesteryl-β-D-galactopyranoside (Table 3). We speculate that membrane insertion could be a possible source of free energy to drive a conformation change of TULIP2 from the ground state structure in Figure 6A and B to a ligand-binding state similar to the structure in Figure 6C. Loops S1-S2 and S4-S5 appear relatively flexible in the molecular dynamics trajectory (Figure 7). In TULIP2, Leucine 105 in loop S4-S5 is predicted to insert into lipid bilayers (Table 2). The free energy change from a water-exposed state to a lipid bilayer-inserted state could spontaneously drive membrane insertion. The locking of loop S4-S5 would, in turn, trigger the separation of α-helix H2 from β strand S1. The interaction between H2 and S1 appears to be one of the weakest tertiary interactions in TULIP2, based on their early separation in our molecular dynamics heating trajectory (Figure 7). Previous models for TULIP conformational changes involved in surfactant properties of latherin ^54^ and SPLUNC1 ^55^ involve a similar flexibility of loops analogous to the TULIP2 S1-S2 loop. In the latherin model, the entire TULIP is proposed to unwrap along a seam analogous to the H2-S1 seam of TULIP2.

The function of TULIP2 in the tick-spirochete-host system remains to be worked out. Our results point to two possibilities: 1) TULIP2 may pick up a lipid molecule from a membrane surface and then transfer it to a receptor on another membrane. For example, *B. burgdorferi* lipids might bind to TULIP2, which then transfers the lipid as a signal to a receptor on a tick immune cell. 2) Alternatively, TULIP2 may function by binding to the *B. burgdorferi* membrane and acting as a scaffold for recruited signaling proteins, thereby recruiting tick hemocytes in an attempt to destroy the spirochetes, by analogy with the bacteriocidal/permeability-increasing protein family of TULIPs.

### 4.2 Purification of TULIPs

The solubility problem we encountered in heterologous expression of TULIP2 might not be unique to TULIP2 but rather common in the TULIP family. A TULIP from *Schistocerca gregaria* locusts, β-carotene-binding protein (BBP), was insoluble when expressed in *E. coli*, but when it was refolded from 8 M urea in the presence of β-carotene BBP was soluble ^56^. Short PLUNC, a single-TULIP domain protein secreted throughout the airways of humans and other mammals, when expressed as a recombinant protein, was reported to precipitate after purification and cleavage of the MBP tag ^57^. Latherin precipitated substantially after tryptic digestion in preparation for mass spectrometry analysis, but solubility issues were not observed in a thioredoxin-tagged form expressed in *E. coli* ^54^. TULIPs involved in cytoplasmic lipid transport (synaptotagmin-like mitochondrial lipid-binding protein, SMP domains) had to be expressed and purified from *E. coli* with MBP tags to enhance solubility for studies of molecular association ^58^.

### 4.3 *I. scapularis* salivary proteome

By comparing saliva proteins from nymphs feeding on *B. burgdorferi*-infected or uninfected mice, we aimed to discover novel protein interactions that contribute to *B. burgdorferi* colonization of the *I. scapularis* midgut. The proteins we identified with expression changes above a significant statistical threshold are shown in Table 1 and Figure 1. Of these proteins, four are lipid-binding proteins (vitellogenin-5, annexin V, saposin, and TULIP2), all of which show decreased levels in saliva of nymphs feeding on *B. burgdorferi*-infected mice; and three are protease inhibitors (trypsin inhibitor, serpin, and chymotrypsin-elastase inhibitor), all of which show increased levels in saliva of nymphs feeding on *B. burgdorferi*-infected mice. A previous study found increased expression of immune-related genes in *B*. *burgdorferi*-infected nymphs ^40^. In our sample, changes in levels of secreted cathepsin L in tick nymphs exposed to *B. burgdorferi* may be immunity-related, as increases in extracellular cathepsins are observed with infections ^59^. Vitellogenin 5 decreased in our *B*. *burgdorferi*-infected nymphs. In the study by Kim et al., *B*. *burgdorferi*-infected nymphs showed a lower expression level of vitellogenin 5, while other vitellogenins, 1, 2, 3 and 6 were up-regulated ^40^. We detected lower amounts of chitinase in nymphs feeding on *B*. *burgdorferi*-infected mice, compared to nymphs feeding on uninfected mice. In contrast, Kim et al. reported an overall higher expression level in infected nymphs. In their study, ticks were infected by force-feeding with BSK-H medium containing *B*. *burgdorferi*, prior to blood feeding, but our ticks had their *B. burgdorferi* exposure in a blood meal as larvae. It is possible that prior spirochete-vertebrate interactions can affect subsequent tick protein levels in response to the spirochete.

One TULIP that we identified in saliva, AEE89467 (Figure 1), was previously characterized as an anticoagulant called p23 ^60^. (The amino acid sequence of p23, accession number HQ605984, is the same as AEE89467). Although the anticoagulant target of p23 was not identified, it is known that the mammalian blood-clotting pathway involves interactions with amino phospholipids on fibroblast and platelet plasma membrane surfaces ^61^, so it is possible that AEE89467 interferes with this interaction through a specific phospholipid binding mechanism.

TULIP2 is a close paralog of AEE89467 (Figure 3). However, docking predictions indicate that TULIP2 is not selective for amino phospholipids over other lipids.

## Acknowledgement

We thank Yue Rachel Chen for help with the early stages of this work.

## Author contributions

Experimental plans: W.O.S, W.P.G., J.S., R.R.

Experimental execution: W.O.S, T.M., W.P.G., J.S., R.R.

Data analysis: W.O.S., W.P.G., J.S., R.R.

Manuscript writing: W.O.S., W.P.G., J.S., R.R.

Computational analysis: M.W.P., L.Y.C., R.R.

