## Supplemental material for "A lipid-binding protein in black-legged tick saliva selectively recognizes *Borrelia burgdorferi* lipids"

**Supplementary Material**

**Figure S1.** Mass spectrometry support for sequences XP_029833768.1 and XP_029824599.1. In 2023, NCBI removed XP_029833768.1 and XP_029824599.1 from their database. XP_029833768.1 was updated to XP_029833768.2, which is now called XP_040079658.2. An NCBI refseq scientist, responding to our inquiry (3-29-23 and 3-30-23), cited new longer reads as the reason for updating these sequences. His opinion was that the sequences, supported by our mass spectrometry data (see MS2 spectrum below), are polymorphisms. Sequences removed by NCBI are shown below with variations highlighted in red.

```
>XP_029833768.1 uncharacterized protein LOC8035502 [Ixodes scapularis]
MLWYGVFAIVLVAYAHAATVQEANHFMDTVLNERLPPLVRASPLLFPVVGIPFFRFDVPKNAPTNRNLHA
NITEGAIRNLDVGVKRMGECLAPALKDGVPTVSCTLDLNNINTTFLAYTKGDNILSTLKEIWVNVLTVDS
VARFEATGNQRRESLLRTFEVPHLHFTTLYNSELHLNTDRQRQFKDHIEAKVKQTLQETLYGDYRLQLAR
AVAATPFPNV

>XP_029824599.1 uncharacterized protein LOC8029408 [Ixodes scapularis]
MLWYGVFAIVLVAYAHAATVQEANHFMDTVLNERLPPLVRASPLLFPVVGIPFFRFDVPKNAPTNRNLHA
NITEGAIRNLDVGVKRMGECLAPALKDGVPTVSCTLDLNNINTTFLAYTKGDNILSTLKEIWVNVLTVDS
IARFEATGVPRRESVLRTFEVPHLHFTTLYNSDLHLNTDRQRQFKDHIETKVKQTLQETLYGDYRLQLAR
AVAATPFPDV

>XP_040079658.2 uncharacterized protein LOC120851145 [Ixodes scapularis]
MLWYGVFAIVLVAYAHAATVQEANHFMDTVLNERLPPLVRTSPLLFPVVGIPFFRFDVPKNAPTNRNLHA
NITEGAIRNLDVGVKRMGECLAPALKDGVPTVSCTLDLNNINTTFLAYTKGDNILSTLKEIWVNVLTVDS
VARFEATGNQRRESLLRTFEVPHLHFTTLYNSELHLNTDRQRQFKDHIEAKVKQTLQETLYGDYRLQLAR
AVAATPFPNV
```

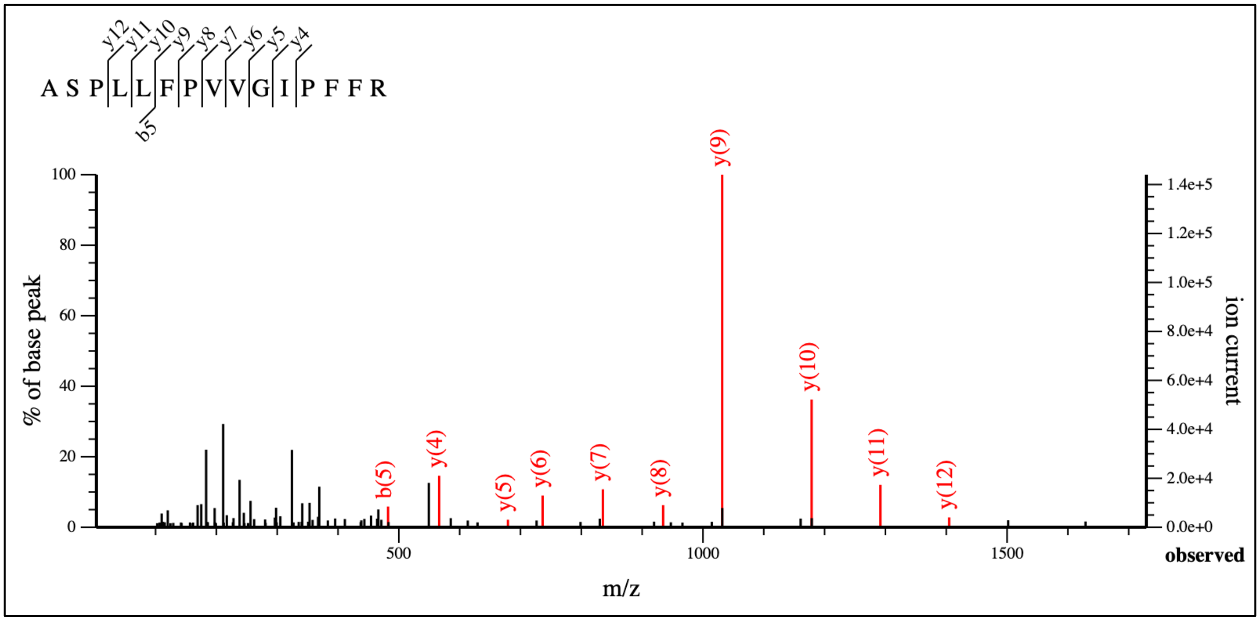

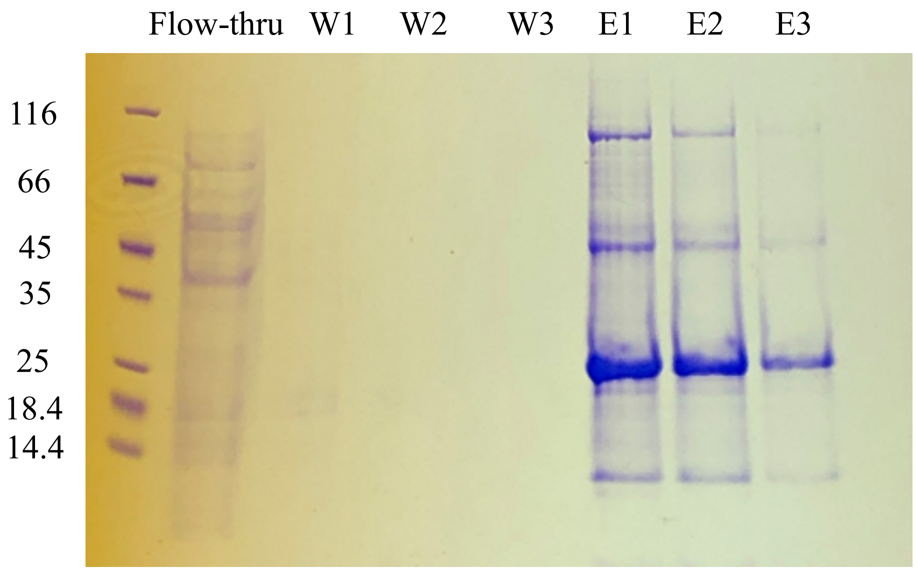


**Figure S2. Coomassie Blue-stained gel image of TULIP2-His purification on Ni-NTA column.** From left to right are molecular weight markers (sizes in kDa labeled on the left), flow-through, three wash fractions (W1 – W3) and three elution fractions (E1 – E3).

**Figure S3. TULIP2 aggregated during dialysis at pH 7.38 and pH 9.34**

Ni affinity-purified TULIP2-His at a concentration of 30 µM, in 5 M Gdn, was dialyzed against PBS (pH 7.38), N-(1,1-Dimethyl-2-hydroxyethyl)-3-amino-2-hydroxypropanesulfonic acid (AMPSO) buffer (pH 9.34) and acetic buffer (pH 5.5; product was not analyzed on the gel). During the dialysis, all setups developed some observable precipitation. The resulting products were centrifuged and analyzed by SDS-PAGE. The supernatant samples did not appear to contain TULIP2-His (~ 25 kDa).

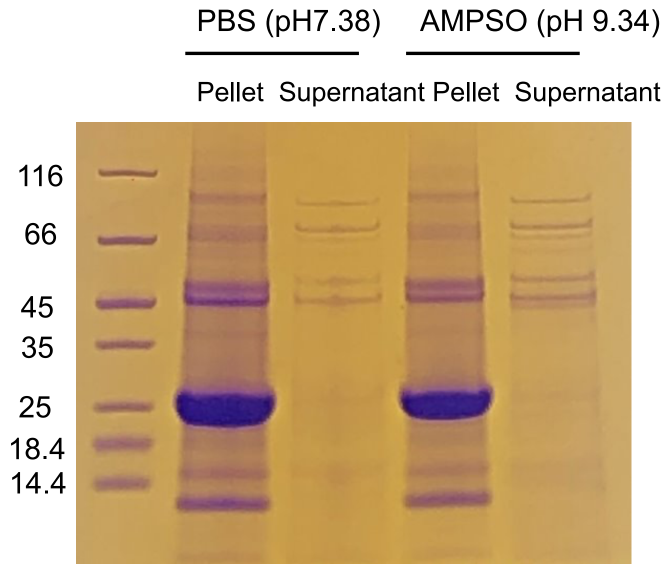


**Figure S3. TULIP2 aggregated during dialysis at pH 7.38 and pH 9.34.**

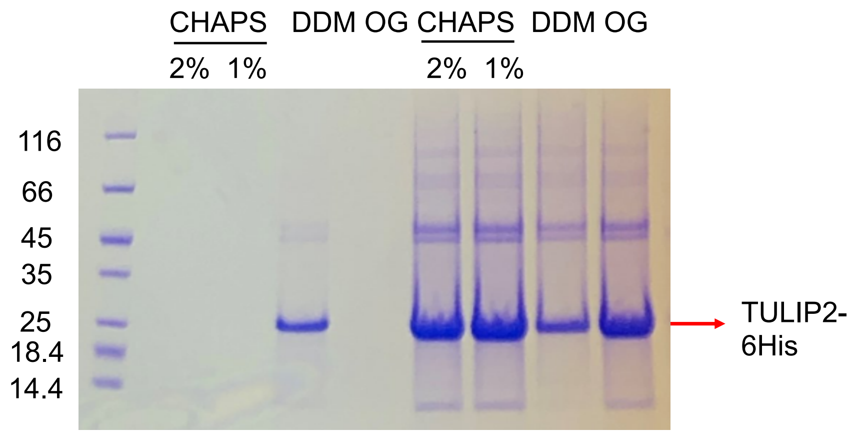


**Figure S4. Effect of three detergents on the aggregation of TULIP2 after dilution to 1 M Gdn.**

Samples loaded from left to right are the supernatant of 2 % CHAPS, 1 % CHAPS, 2% dodecyl maltoside (DDM) and 1.5 % octyl glucoside (OG), and the pellet suspension in the same order.

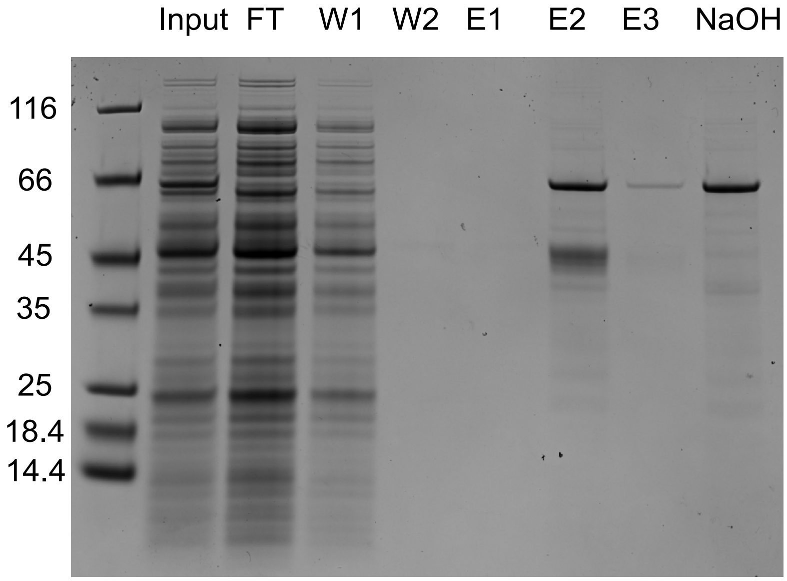


**Figure S5. Purification of MBP-TULIP2 on MBPTrap column.** *E. coli* lysate (input) was applied to MBPTrap column, and the flow-through (FT) was collected. The column was washed with 10-bed volume of binding buffer. The first one ml (W1) and the last one ml (W2) was examined by gel electrophoresis. Three one-ml elution fractions (E1-E3) and the first one ml of NaOH regeneration were examined as well.

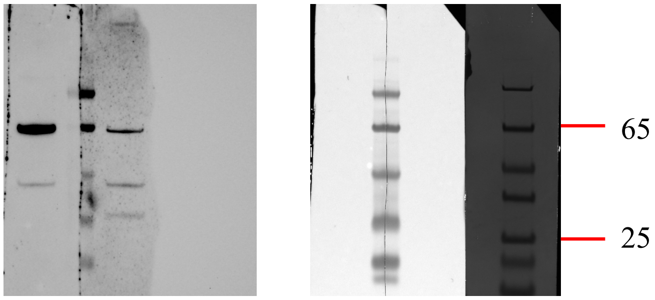


**Figure S6. Western blot of MBP-TULIP2.** Photo on the left is chemiluminescence, and colorimetric on the right. Lanes 1 and 3 are the same MBP-TULIP2 sample, Lane 2 is Thermo Fisher Pierce prestained marker and the last lane is Pierce unstained marker. The positions of 65 and 25 kDa bands are marked according to the unstained marker. In the chemiluminescence photo, the left blot was probed with anti-MBP primary antibody, and the right blot was probed with anti-TULIP2 antiserum.

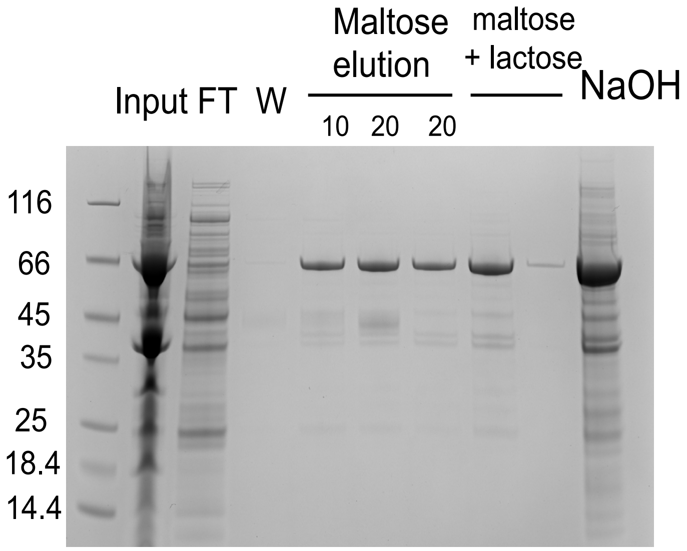


**Figure S7. Lactose washed out some MBP-TULIP2 that did not elute in 20 mM maltose.** Maltose-affinity column purification fractions were analyzed on SDS-PAGE gel in the loading order of column input, flow-through (FT), final wash fraction (W), three elution fractions with indicated concentrations (mM) of maltose, two elution fractions of 20 mM maltose plus 300 mM lactose and NaOH regenerative wash.

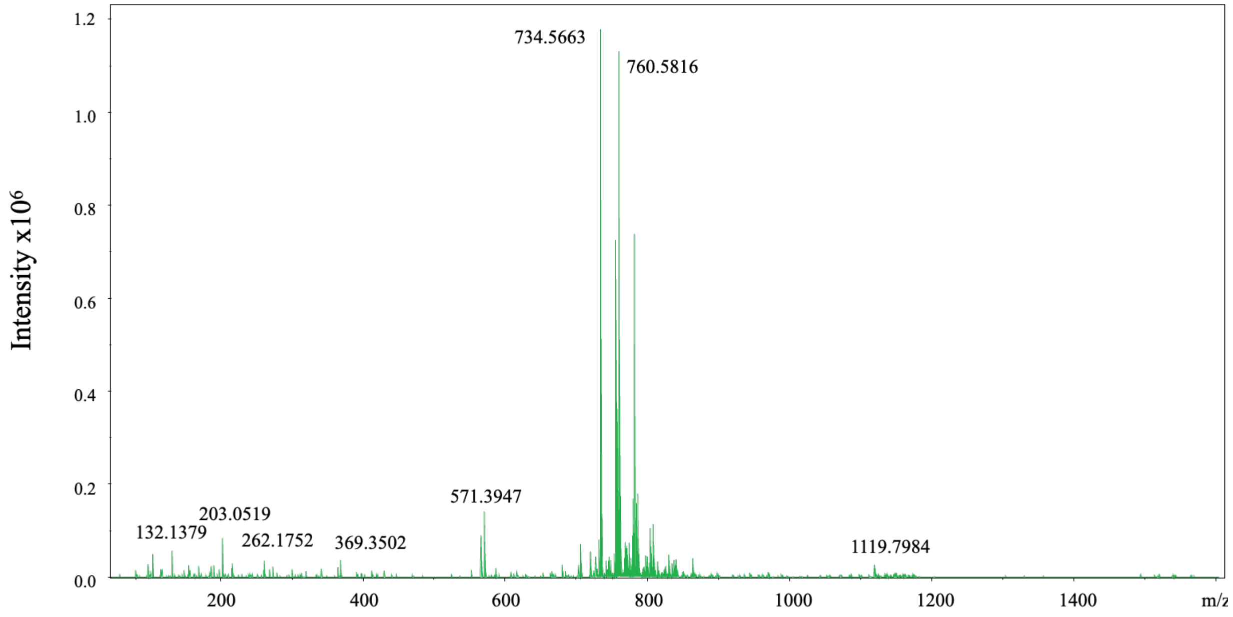


**Figure S8. ESI-QTOF mass spectrum of *B. burgdorferi* lipid extract.**

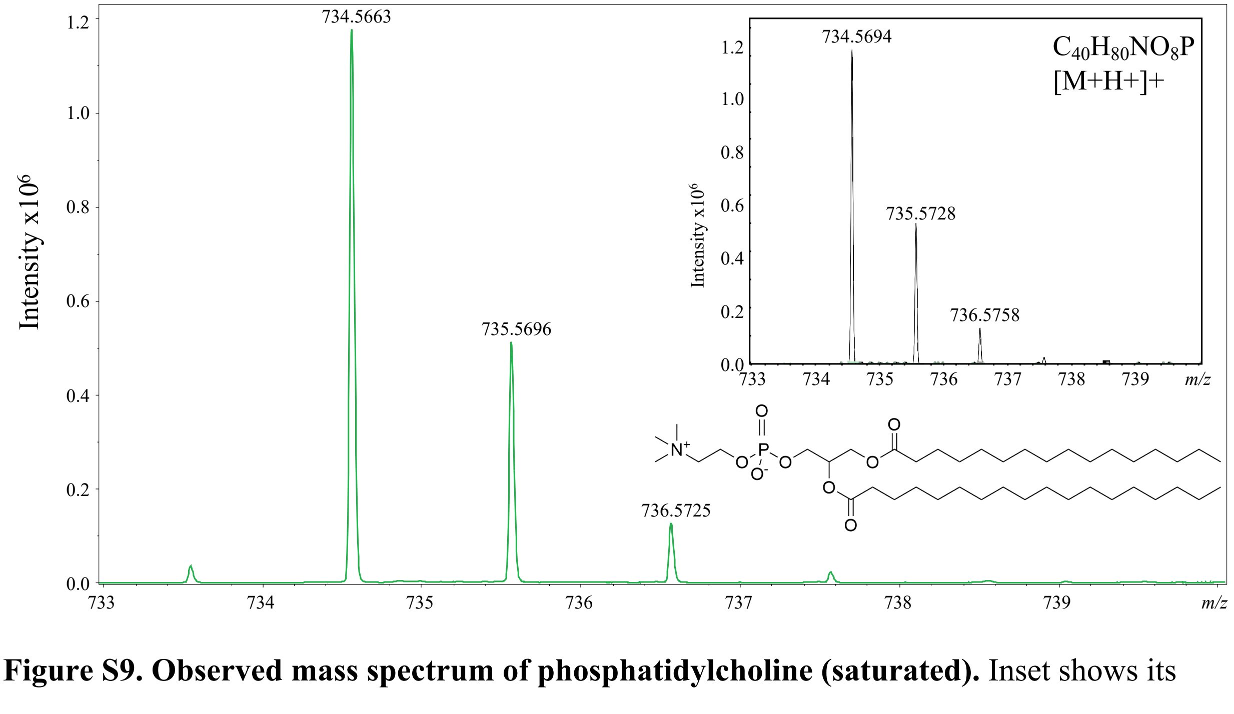


**Figure S9. Observed mass spectrum of phosphatidylcholine (saturated).** Inset shows its theoretical spectrum and structural representation.

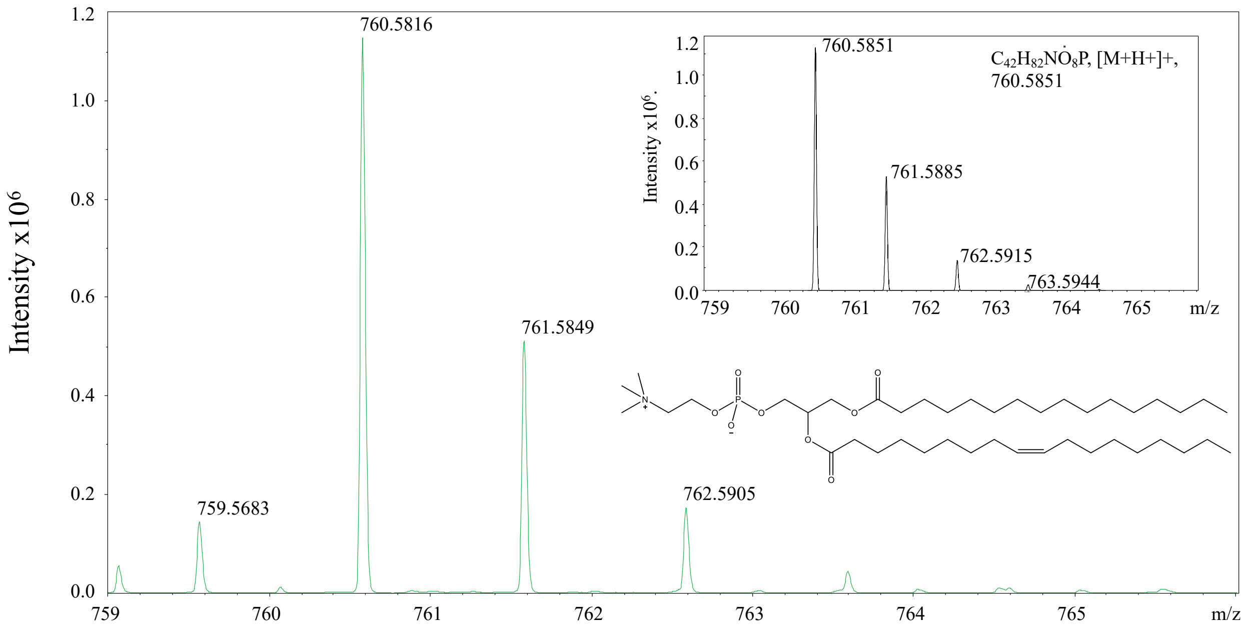


**Figure S10. Observed mass spectrum of phosphatidylcholine with one double bond.** Inset shows its theoretical spectrum and structural representation.

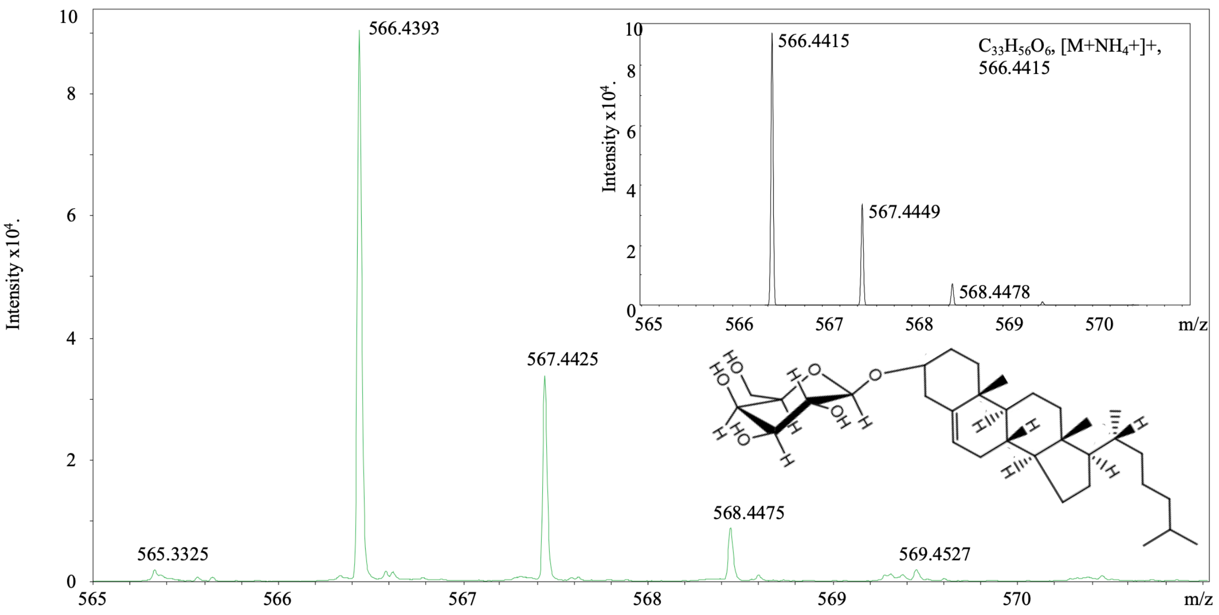


**Figure S11. Observed mass spectrum of cholesteryl-β-D-galacto-pyranoside.** Inset shows its theoretical spectrum and structural representation.

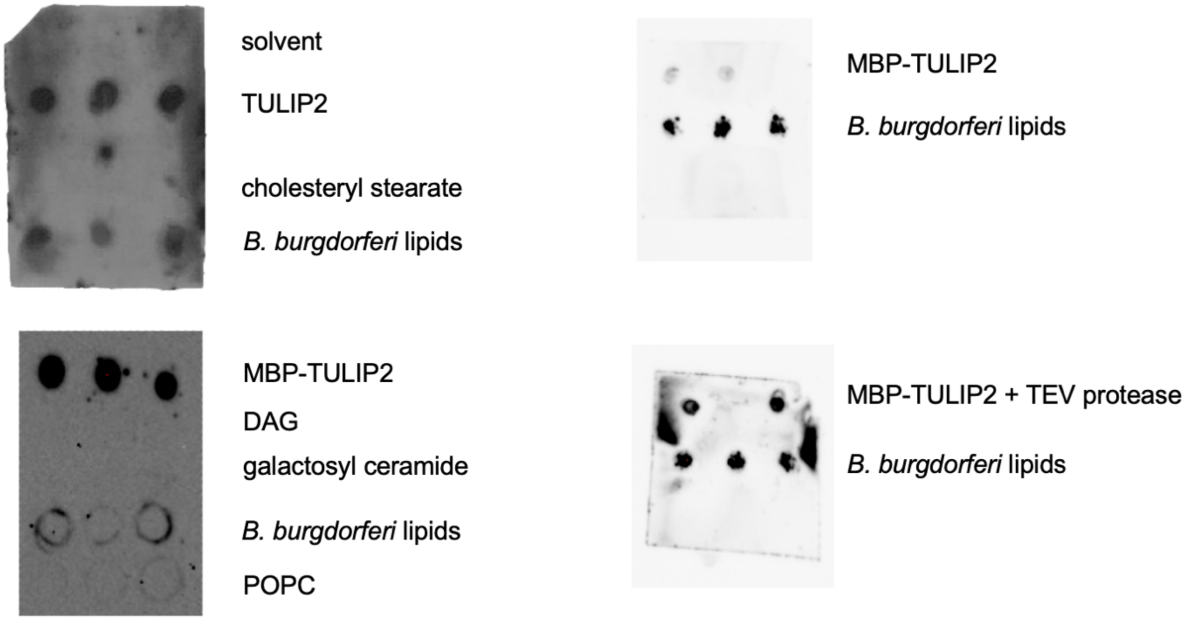


**Figure S12. Protein-lipid overlay assay.** Extracted *B. burgdorferi* lipids and control lipids were spotted on nitrocellulose film (see Figure 5). The films were blocked and then equilibrated with mouse anti-TULIP2 serum, followed by horseradish peroxidase-linked anti-mouse IgG, and imaged by chemiluminescence. The presence of the MBP tag did not interfere with binding to *B. burgdorferi* lipids (right panel).

**Table S1. Aggregation of TULIP2 after dialysis in the presence of mild detergents at the indicated concentrations.**

| | PBS | 2% CHAPS | 1% CHAPS | 1.5% OG | 2% DDM |
|---|---|---|---|---|---|
| A330 turbidity | 1.47 | 2.24 | 2.10 | 2.18 | 0.45 |
| A280 | 0 | 0.033 | 0.003 | 0.159 | 0.182 |
